# Site-Specific Synthesis of N4-Acetylcytidine in RNA Reveals Physiological Duplex Stabilization

**DOI:** 10.1101/2021.11.12.468326

**Authors:** David Bartee, Kellie D. Nance, Jordan L. Meier

## Abstract

*N*^4^-acetylcytidine (ac4C) is a post-transcriptional modification of RNA that is conserved across all domains of life. All characterized sites of ac4C in eukaryotic RNA occur in the central nucleotide of a 5’-CCG-3’ consensus sequence. However, the thermodynamic consequences of cytidine acetylation in this context have never been assessed due to its challenging synthesis. Here we report the synthesis and biophysical characterization of ac4C in its endogenous eukaryotic sequence context. First, we develop a synthetic route to homogenous RNAs containing electrophilic acetyl groups. Next, we use thermal denaturation to interrogate the effects of ac4C on duplex stability and mismatch discrimination in a native sequence found in human ribosomal RNA. Finally, we demonstrate the ability of this chemistry to incorporate ac4C into the complex modification landscape of human tRNA, and use duplex melting combined with sequence analysis to highlight a potentially unique enforcing role for ac4C in this setting. By enabling the analysis of nucleic acid acetylation in its physiological sequence context, these studies establish a chemical foundation for understanding the function of a universally-conserved nucleobase in biology and disease.

## Introduction

*N*^4^-acetylcytidine (ac4C) is a modified RNA nucleobase that is universally conserved amongst all domains of life (Figure 1a).^1^ Cytidine acetylation was first identified in eukaryotic transfer RNA (tRNAs) in the 1960’s.^2,3^ Subsequent quantitative mapping studies have defined helices 34 and 45 of 18S ribosomal RNA (rRNA) and the D-stem of tRNA^Ser^ and tRNA^Leu^ as the dominant sites of ac4C in eukaryotes.^4–6^ In humans, cytidine acetylation is catalyzed by the essential RNA acetyltransferase enzyme NAT10, which works in concert with protein and snoRNA adapters to address its distinct targets.^4,5^ Dysregulation of NAT10 has been associated with many diseases, including premature aging syndromes and cancer.^7,8^ Precisely why ac4C is so highly conserved in eukaryotic rRNA and tRNA remains unknown.

**Figure 1.**
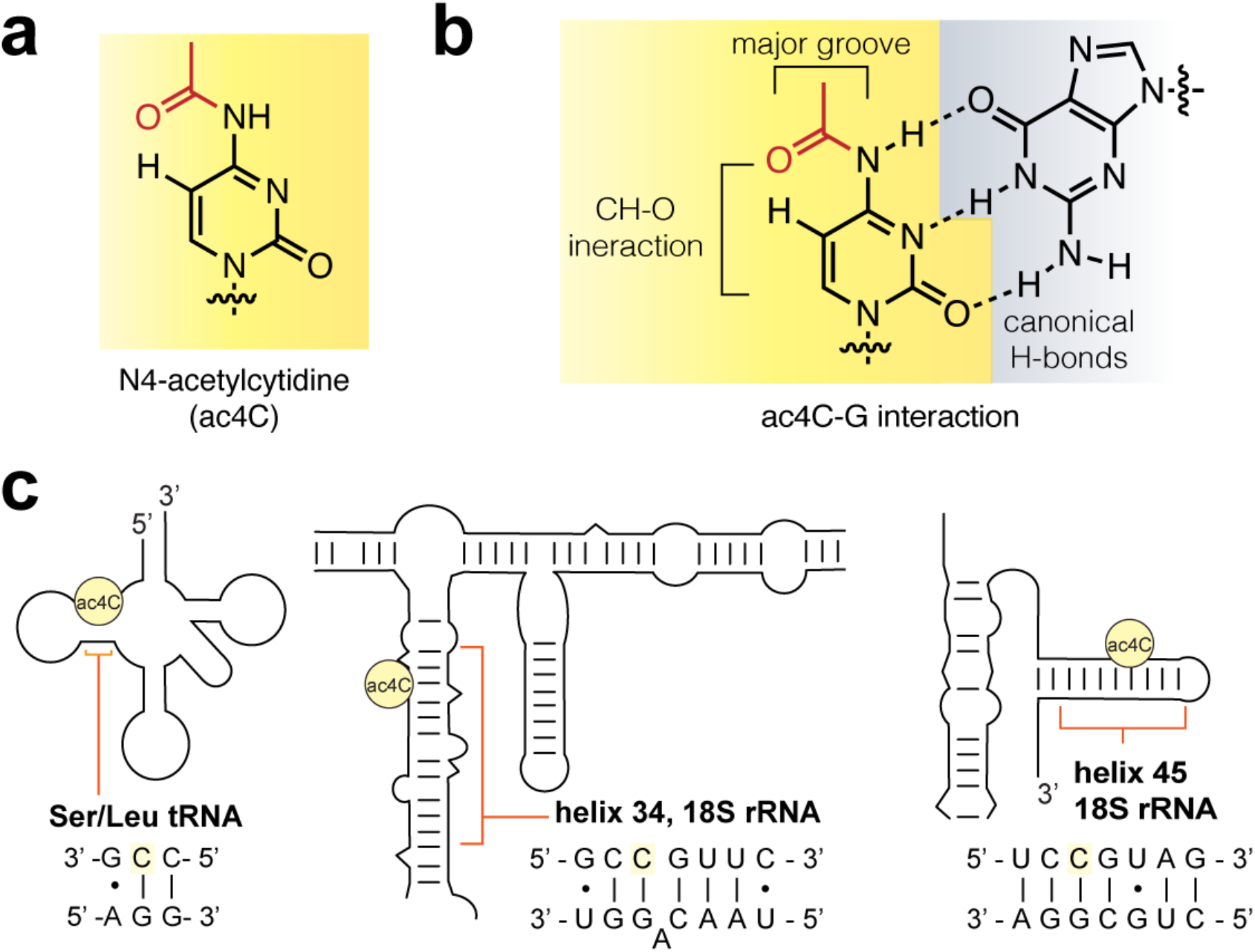
(a) N4-acetylcytidine (ac4C). (b) Schematic of ac4C-G base pair. (c) Sites of ac4C that have been identified using nucleotide resolution methods in human RNA.

Knowledge of the molecular effects of ac4C largely derive from modeling and structural studies of the free nucleoside.^9,10^ Crystallographic data indicate the *N*^4^-acetyl group in ac4C prefers a conformation in which it is oriented proximal to cytidine C^5^, reflecting the influence of a weak C-H···O interaction formed between the acetamide carbonyl oxygel and pyrimidine C^5^ C-H (Figure 1b).*^15, 16^* This ordered structure is compatible with canonical base-pairing, as it places the bulk of the acetyl group towards the major groove of duplex RNA. *N*^4^-acetylation also stabilizes the C3’-endo conformation of cytidine’s ribose sugar, an effect common to other rRNA modifications such as pseudouridine and 2’-*O*-methylation.^11^ These features were recently corroborated in a series of high resolution cryo-EM structures of eukaryotic and archaeal ribosomes.^6,12,13^

Every specific site of ac4C thus far localized in human RNA occurs at the central base of a 5’-C**C**G-3’ consensus sequence (Figure 1c).^14^ An identical 5’-C**C**G-3’ sequence is acetylated in members of the archaeal order Thermococcales, whose RNA harbors the most ac4C of any organism yet characterized on Earth.^15,16^ Interestingly, many cytidines that are dynamically acetylated in response to temperature in Thermococcales occur at the stem of hairpin structures adjacent the loop, suggestive of a role for ac4C in enforcing duplex stability.^6^ Biophysical analyses of ac4C would provide foundational data as to its role in biology and disease. However, ac4C has yet to be characterized in any physiologically-relevant sequence context due to a lack of methods to site-specifically introduce it into RNA.

Previous studies have incorporated ac4C into RNA enzymatically using in vitro transcription.^17,18^ While this approach facilitates many applications, it results in a non-physiological, homogenous replacement of every templated cytidine with an ac4C. Conventional protocols for solid-phase synthesis of RNA oligonucleotides are similarly incapable of producing ac4C RNA.^19^ This is because these methods employ *N*^4^-acetylation to protect the exocyclic amine of cytidine during iterative coupling and deprotection steps, and thus have been designed (even in the case of “fast-deprotecting” phenoxyacyl protection)^20^ to remove this modification during nucleobase deprotection or upon nucleophilic cleavage from ester-linked resins (Figure 2a). Despite being listed as a potential component of nucleic acid therapeutics in many patent applications,^21^ the synthesis and characterization of ac4C at defined positions in RNA has never been reported. The synthesis of site-specific acetylated RNA is a prerequisite for understanding the biological role of ac4C and applying it as a functional element in nucleic acid therapeutics. These opportunities highlight the need for a synthetic route.

**Figure 2.**
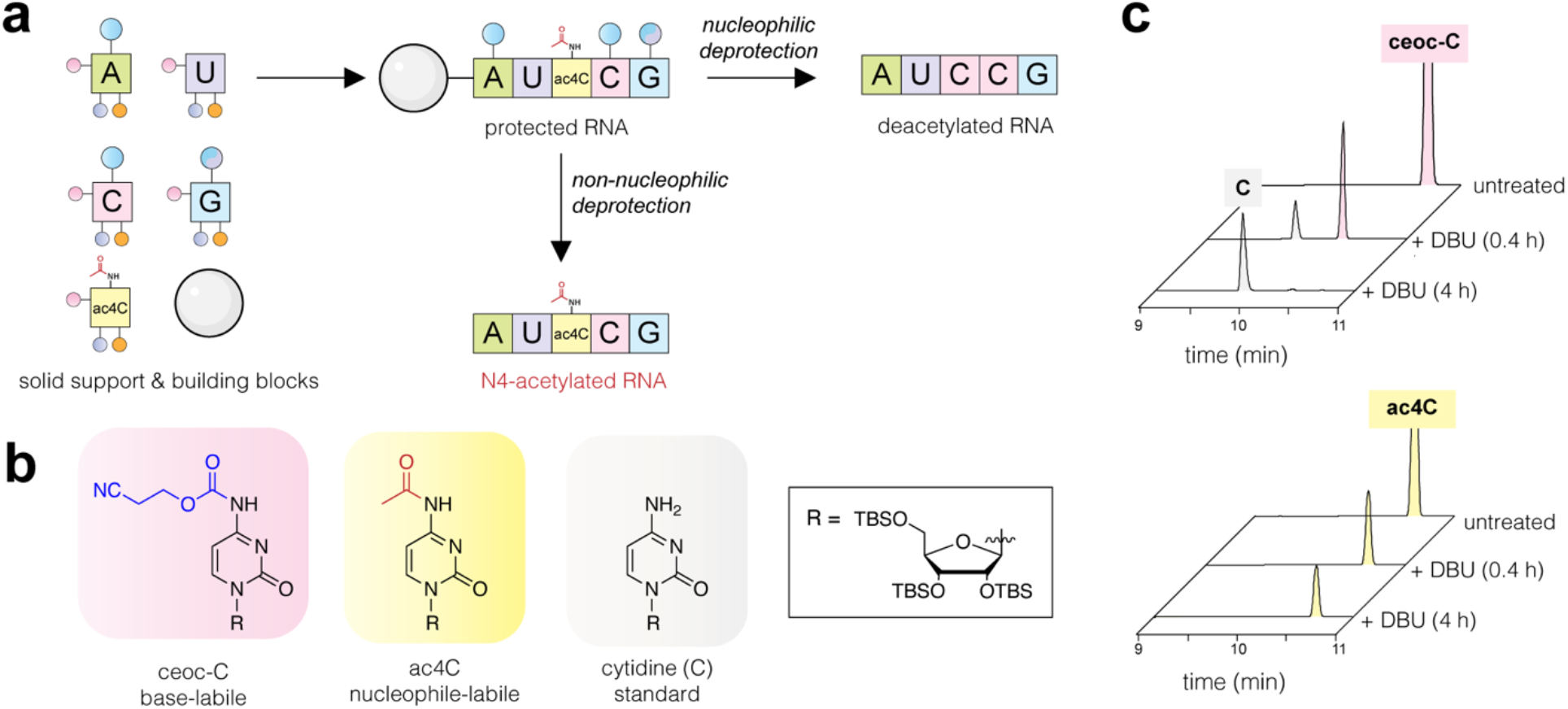
(a) Strategy for site-specific synthesis of ac4C RNA. Retention of ac4C requires a solid support and protected building blocks that are labile to non-nucleophilic condition (center), thus avoiding nucleophilic deprotection which cleaves ac4C (right). (b) Model substrates used in ac4C-DBU compatibility studies. (c) HPLC traces of model ceoc-C (top) and ac4C (bottom) substrates following exposure to DBU for 0.4 or 4 h. Extended HPLC traces are provided in the Supplementary Information.

## Results and Discussion

### An orthogonal protection strategy compatible with cytidine acetylation

Site-specific incorporation of cytidine acetylation into RNA requires i) a protecting group for the exocyclic nucleobase nitrogens and ii) a solid-phase support linkage that can be cleaved without removing the *N*^4^-acetyl group of ac4C (Figure 2a). To address the first criterion, we were inspired by prior syntheses of *O*-acetylated RNAs, which like ac4C are sensitive to nucleophilic cleavage.^22,23^ These studies employed *N*-cyanoethyl *O*-carbamate (*N-*ceoc)^24^ nucleobases that could be deprotected using the non-nucleophilic base 1,5-diazabicyclo(4.3.0)non-5-ene (DBU). To determine the orthogonality of *N*-ceoc protection and *N*^4^-acetylation, we analyzed the compatibility of DBU and ac4C using a series of simple model substrates (Figure 2b). Over 4h, DBU cleanly removed the *N*-ceoc group from ceoc-C, while leaving *N*^4^-acetylation intact (Figure 2c). However, degradation was observed in the presence of morpholine (10% v/v) (Figure S1). The exquisite sensitivity of ac4C to nucleophilic cleavage reagents differentiates this study from prior work, which were able to use morpholine to scavenge acrylonitrile during the synthesis of *O*-acetylated RNAs.^22^ These studies define an orthogonal condition for the protection and deprotection of ac4C-containing RNA.

### Synthesis of building blocks and solid-phase for site-specific ac4C RNA synthesis

To develop our strategy in an oligonucleotide context, we next synthesized *N-*ceoc protected phosphoramidites of adenosine, cytidine, and guanosine via 3’,5’-cyclic silyl protected strategy (Figure 3, top).^25^ Briefly, parent nucleosides were first protected at the ribose sugar via treatment with di-tertbutylsilyl bistriflate, followed by addition of tert-butyl dimethylsilyl chloride and imidazole. In the case of cytidine, the pyrimidine ring was protonated using one equivalent of triflic acid prior to addition of the silyl bistriflate. After protection of ribose, the exocyclic nitrogens were carbamoylated using ceoc-carbonyl-*N*-methylimidazolium chloride. In the case of guanosine, the *O*^6^ was first protected via Mitsunobu reaction with (4-nitrophenyl) ethanol prior to carbamoylation at *N*^2^ using ceoc-chloroformate. Selective removal of the 3’,5’-cyclic silyl ether was achieved using HF-pyridine in dichloromethane. Finally, regioselective introduction of dimethoxy at the 5’ position, followed by phosphitylation, yielded A, C, and G phosphoramidite monomers in sufficient yields for solid-phase synthesis.

**Figure 3.**
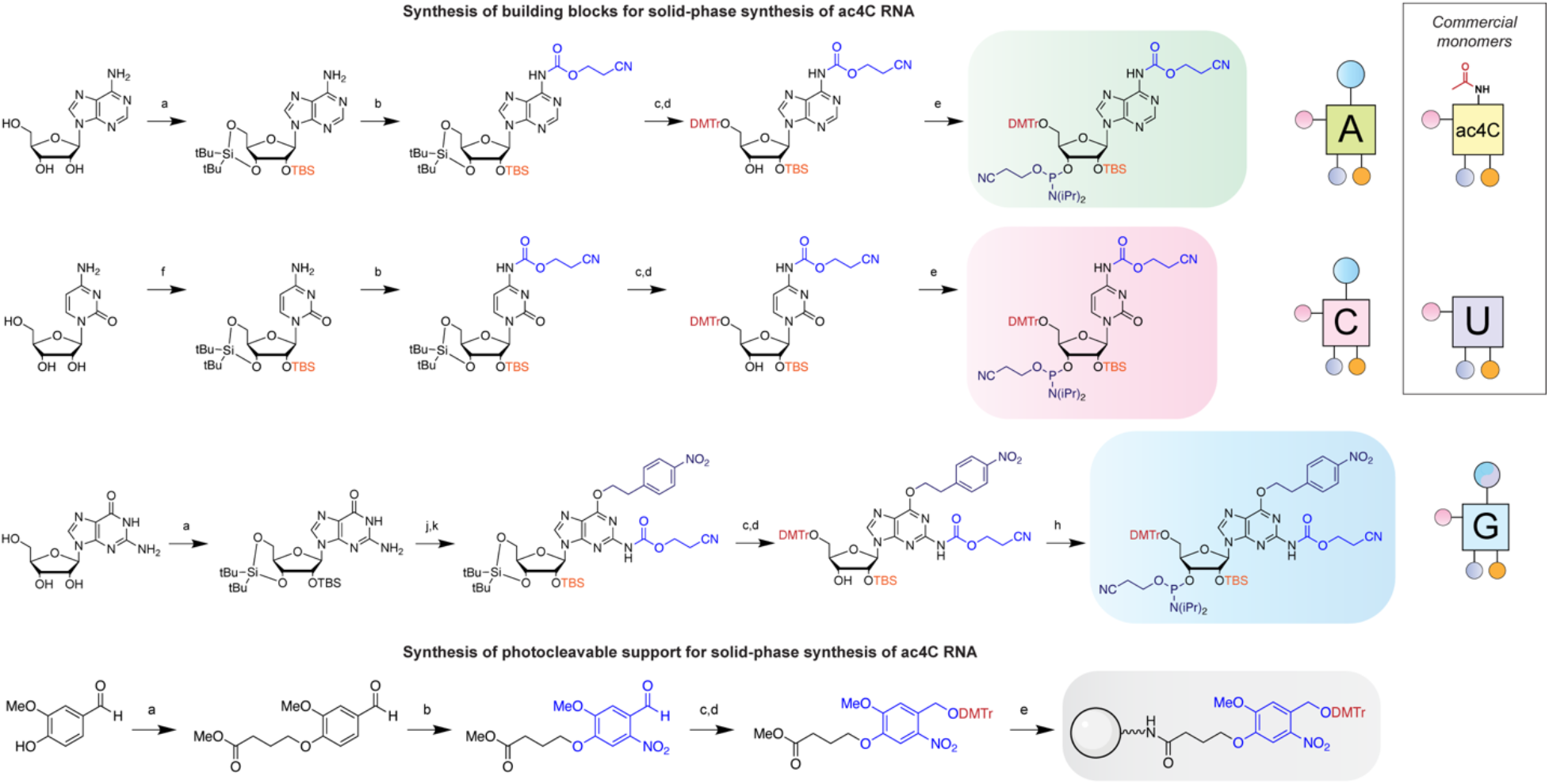
Synthesis of building blocks for site-specific ac4C synthesis. Top: synthesis of *N*-ceoc-protected phosphoramidites. a) i) (O-tBu)_2_Si(OTf)_2_, DMF, 0 °C, ii) TBS-Cl, imidazole, DMF, 60 °C; b) ceoc-carbonyl-*N*-methylimidazolium chloride, DCM, 23 °C; c) HF-pyridine, pyridine, DCM, 0 °C; d) DMTr-Cl, pyridine, 4 °C; e) (OCH_2_CH_2_CN)P(iPr2N)_2_, tetrazole, ACN, 23 °C; f) (O-tBu)_2_Si(OTf)_2_, TfOH, DMF, 0 °C, ii) TBS-Cl, imidazole, DMF, 60 °C; g) H_2_ (1 atm), Rh/alumina, MeOH, 23 °C; h) (OCH_2_CH_2_CN)(iPr_2_N)PCl, iPr_2_NEt, THF, 23 °C; j) triphenylphosphine, DIAD, (4-nitrophenyl)ethanol, dioxane, 100 °C; k) ceoc-chloroformate, DCM, 23 °C. Figure 4. Bottom: synthesis of photocleavable solid support. a) Methyl (4-bromobutryate), K_z_CO_3_, DMF, 23 °C; b) TFA, KNO_3_, THF, 23 °C; c) NaBH_4_, MeOH, 23 °C, d) DMTr-Cl, pyridine, 4 °C; e) i) aq. LiOH, THF, 23 °C, ii) HATU, iPr_2_NEt, LCAA-CPG, ACN, 23 °C, iii) Piv-Cl, *N*-methylimidazole, 2,6-lutidine, THF, 23 °C. Right: graphical abbreviations for monomers used in this study. Full NMR and mass spectra are provided in the Supplementary Information.

Next, we sought to devise a solid-phase support that could release RNA oligonucleotides without deacetylating ac4C. Given the incompatibility of ac4C with nucleophiles, we chose to pursue a photocleavable approach. We hypothesized that a nitroveratryl-based linker may be optimal for this purpose, allowing for mild cleavage upon irradiation at 365 nm while minimizing photochemical reactions of RNA caused by lower wavelength UV light (Figure 3, bottom). This necessity is underscored by ac4C’s relatively red-shifted absorbance (λ_max_ = 302 nm) and previously observed photochemistry.^26,27^ Synthesis of the linker began with alkylation of vanillin followed by trifluoroacetic acid-mediated nitration. Reduction of the aldehyde afforded nitroveratryl alcohol, which was further protected with dimethoxytrityl chloride in pyridine to yield the elaborated linker. Subsequent deprotection and coupling to long-chain alkylamine derivatized controlled pore glass (LCAA-CPG) provided access to photocleavable solid-support.

### Synthetic optimization enables site-specific incorporation of ac4C in RNA

With these reagents in hand, we set out to establish conditions for the synthesis of ac4C RNA oligomers (Figure 3a). These studies employed standard phosphoramidite coupling time (6 min), coupling reagents (ETT), oxidation conditions (I_2_, pyridine, H_2_O), and decapping reagent (3% TCA in DCM). To avoid reaction of acetic anhydride with *N*-protected exocyclic amines conventional 5’-OH capping was omitted. This step was further determined to be dispensable based on production of similar amounts of full-length RNA in uncapped and pivalic anhydride-capped samples (Figure S2).^28^ However, while solid-phase synthesis proved straightforward, successfully isolating homogenous *N*^4^-acetylated RNA required several innovations compared to previous approaches. First, to avoid nucleobase alkylation by acrylonitrile during *N*-ceoc removal we developed an on-column deprotection scheme (Figure 4a, optimization #1). This protocol passages DBU (0.5 M in acetonitrile) over the nascent RNA oligomer on solid-support to efficiently deprotect *N*-ceoc bases while limiting their exposure to acrylonitrile thus obviating the need for a nucleophilic scavenger such as morpholine. Second, to maximize yields of ac4C-containing RNA, we identified photolysis conditions that efficiently cleave product from solid support but minimize unintentional *N*^4^-deacetylation (Figure 4a, optimization #2). Initial experiments using a model RNA (5’-UU(ac4C)UUp-3’) indicated ~47% of ac4C was deacetylated during photolysis or 2’-*O*-TBS removal (Figure S3). Addition of Hunig’s base to the desilylation reaction modestly impeded deacetylation (47% to 35%, optimization #3). Changing the photolysis solvent to buffered acetonitrile had a more profound effect, reducing the extent of deacetylation to less than 5% (Figure S3). Elimination of these deacetylation products greatly facilitates the synthesis of ac4C-containing RNAs by both improving yield and streamlining purification. Finally, inspired by the work of Sekine et al.,^29^ we tested the synthesis using an *N*-unprotected guanosine phosphoramidite (Figure 4b). The use of this synthetically facile monomer further improved accumulation of full-length RNA products. Combining these innovations resulted in crude cleavage reactions that contain high proportions of the desired products (Figure S4, Figure 4c) which could be further purified using polyacrylamide electrophoresis (PAGE) to yield pure ac4C-containing RNAs (Figure 4d). Overall, these studies define an effective solid-phase synthetic route to RNAs containing the endogenous electrophilic base modification ac4C.

**Figure 4.**
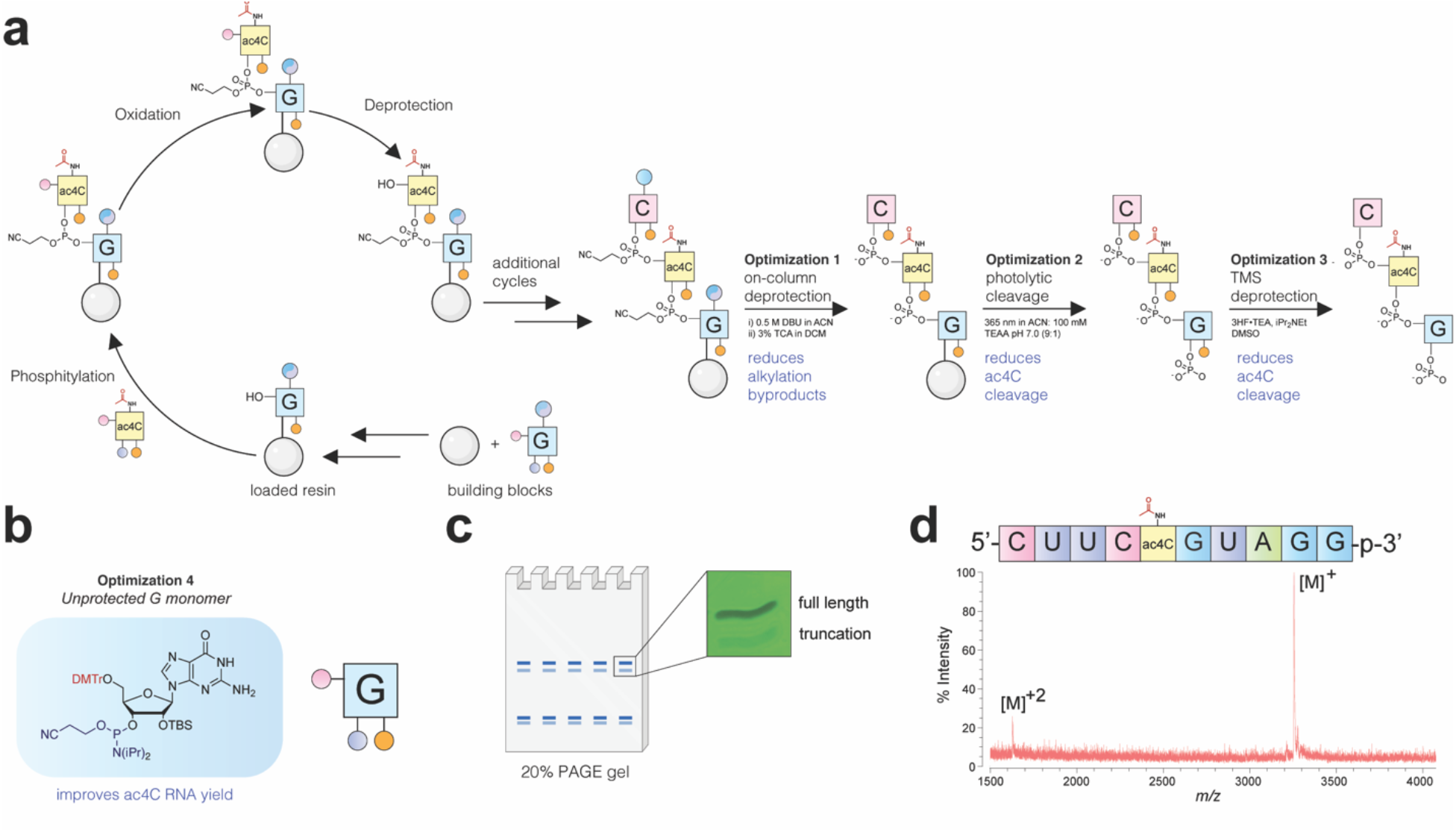
(a) Solid-phase synthesis of ac4C-containing RNA oligonucleotides. Standard phosphoramidite synthesis conditions were employed, with the exception that no 5’-capping step was used. Optimizations critical for ac4C synthesis included on-column deprotection (#1), buffered photolytic cleavage (#2), and buffered desilylation condition (#3) to minimize alkylation and ac4C cleavage byproducts during release of the deprotected oligomer. (b) Ac4C synthesis is improved by the use of unprotected G monomer (#4). (c) Schematic for polyacrylamide gel electrophoresis (PAGE) purification and UV image of full-length and truncation products formed by optimized synthesis. (d) MALDI-TOF mass spectra of purified ac4C-containing 10-mer RNA. Full gels and MALDI-TOF spectra are provided in the Supplementary Information.

### Synthesis and characterization of ac4C in an endogenous 5’-CCG-3’ sequence context: 18S rRNA

Human small subunit (SSU) 18S rRNA contains two high stoichiometry sites of ac4C (C1280 and C1842), each of which is embedded in a fully base-paired 5’-CCG-3’ sequence (Figure 1b-c).^4–6^ To study cytidine acetylation in this context, we synthesized an RNA decamer corresponding to the ac4C-containing strand of SSU helix 45 (Table 1). Annealing to a complementary RNA enabled analysis of ac4C’s effects on duplex stability and mismatch discrimination via UV-melting experiments. Thermal denaturation curves were analyzed by both non-linear regression and van’t Hoff plots to extract thermodynamic constants (Table 1, Supplementary Information). Agreement between these two analysis methods supports a two-state denaturation model for all duplexes analyzed. Focusing first on a fully complementary duplex (Table 1, entry 1), cytidine acetylation was found to have an overall stabilizing effect on duplex RNA (ΔTm_ac4C v. C_ = +1.7 °C). The free energy change caused by ac4C is accounted for by increased enthalpy upon duplex formation relative to cytidine RNA. The only previous study of *N*^4^-acetylcytidine in a hybridized oligonucleotide context was in a polyuridine DNA, and observed a smaller increase in melting temperature (ΔTm_ac4C v. C_ = +0.4 °C).^30^ Further study will be required to determine whether this difference reflects unique experimental conditions or a stabilizing effect of ac4C on its evolutionarily conserved RNA sequence context. Duplexes containing mismatches across from cytidine or ac4C mismatch duplexes each exhibited reduced melting temperatures relative to their match counterparts (Table 1, entries 2-4). Overall, the effects of ac4C on mismatch discrimination (ΔΔTm_C-G v. C-A_) are small and within the error of our experimental measurement. Taking into account the average stabilities of match and mismatch, substitution of cytidine with ac4C appears to slightly discriminate against a C-A mismatch (ΔΔTm = −0.4 °C) and increase tolerance for C-U mismatch (ΔΔTm = +2.0 °C). Improved C-A mismatch discrimination by ac4C is consistent with prior studies of *E. coli* tRNA^Met^, where incorporation of N^4^-acetylation has been shown to prevent misreading of AUA codons.^31,32^ The potential for ac4C to engage in non-canonical pairing with uridine has not been previously described, but is at least anecdotally supported by the recent cryo-EM visualization of this base pair in an archaeal ribosome (Figure S5).^6^

**Table 1.**
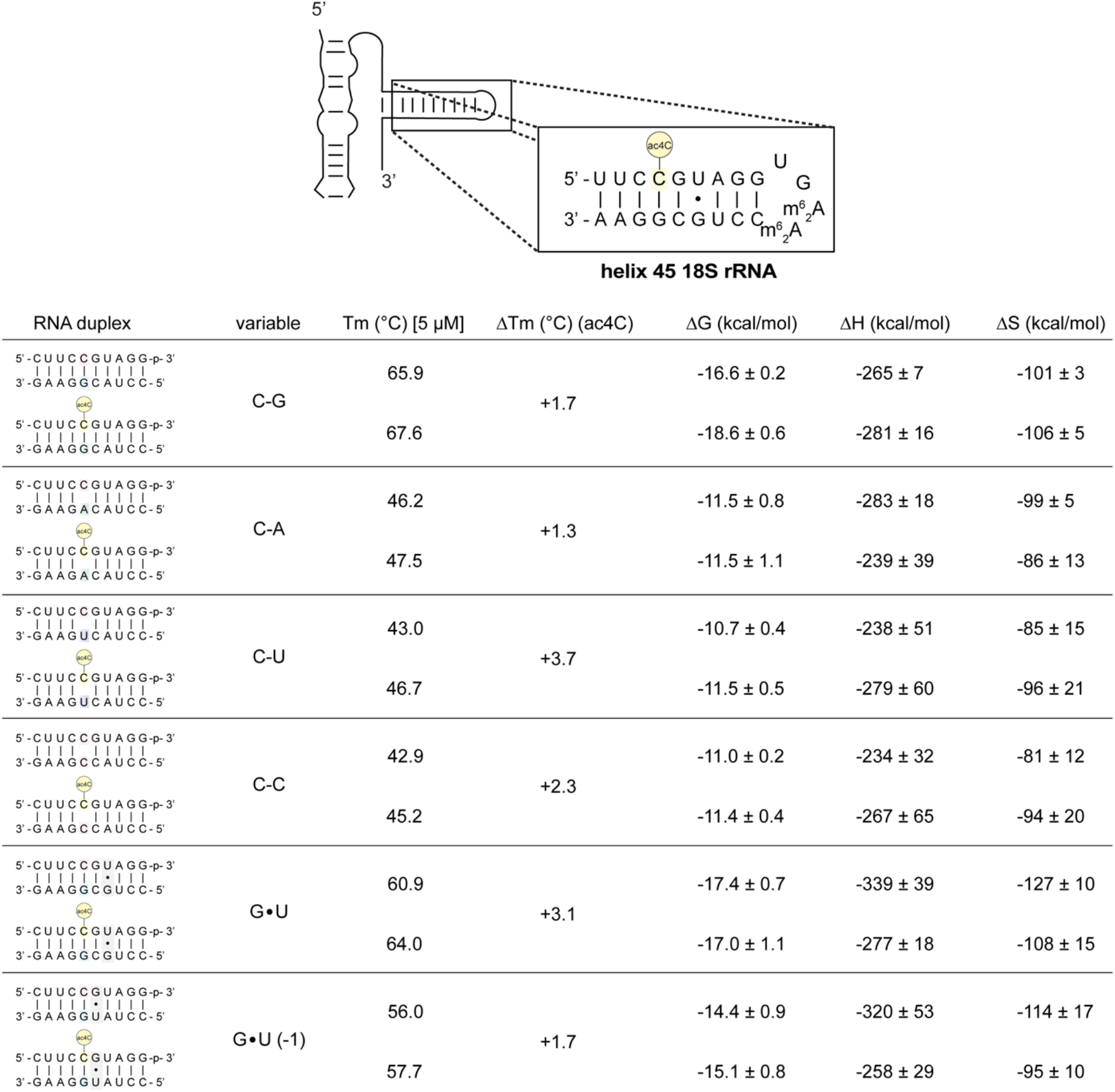
UV-melting of RNA duplexes inspired by human SSU rRNA helix 45 (top). Duplexes were designed to test the effect of ac4C on canonical base-pairing, mismatch discrimination, and compatibility with adjacent G●U wobble pairs (n=3). ΔTm = Tmac4C v TmC [5 μM]. Exemplary melting curves and van’t Hoff plots are provided in the Supplementary Information.

Both known sites of cytidine acetylation in human rRNA reside two bases from a G●U wobble base pair (Figure 1b-c).^4–6^ Given the significance of G●U pairing to RNA structure,^33^ we next set out to determine how ac4C alters the stability of duplex RNAs containing this element. Cytidine and ac4C duplexes were prepared containing a G●U pair +2 bp from ac4C, effectively replicating the stem sequence found in helix 45 of human 18S rRNA. Once again, RNA duplexes containing ac4C were found to be more stable than those containing cytidine (ΔTm_ac4C v. C_ = +3.1 °C, entry 5). Cytidine acetylation also stabilizes duplexes containing a G●U pair directly proximal to the modified nucleotide (entry 6), albeit to a lesser extent.

Differences in basal stability confound a quantitative comparison of G●U versus and fully complementary RNA duplexes (entries 1 and 5). However, the observation that ac4C is slightly more stabilizing in the G●U duplex (3.1 °C vs. 1.7 °C) indicates the high compatibility of this modification with adjacent non-canonical RNA base pairs. Moreover, these studies provide the first empirical evidence that site-specific cytidine acetylation can enforce RNA structure in a physiologically-relevant sequence context.

### Synthesis of ac4C in a complex modification landscape: tRNA^Ser^

Eukaryotic tRNA^Ser^ constitutes the first site of cytidine acetylation ever characterized.^2,3^ Deposition of ac4C in eukaryotic tRNAs occurs at C12 of the D-arm, is exclusive to tRNA^Ser/Leu^ (Figure 5a), and requires both a cytidine acetyltransferase (Nat10 in humans; Kre33 in yeast) <u>and</u> an adapter protein (Thumpd1 in humans; Tan1 in yeast).^5,34^ Deletion of yeast Tan1 causes loss of tRNA^Ser^, rapid decay of tRNA^Ser^_AGA_, and growth defects at elevated temperatures.^35^ This could indicate a critical role for ac4C in enforcing tRNA^Ser^ structure or, alternatively, reflect an ac4C-independent effect caused by loss of Tan1. Emphasizing the need to consider this latter possibility, previous studies have found proteins that carry out tRNA modifications can aid tRNA maturation independent of their catalytic activity.^36^ Differentiating between these scenarios would be greatly aided by the ability to isolate the biophysical effects of ac4C in the unique context of the tRNA D-arm. This led us to ask the question: does cytidine acetylation alter the stability of tRNA^Ser^?

**Figure 5.**
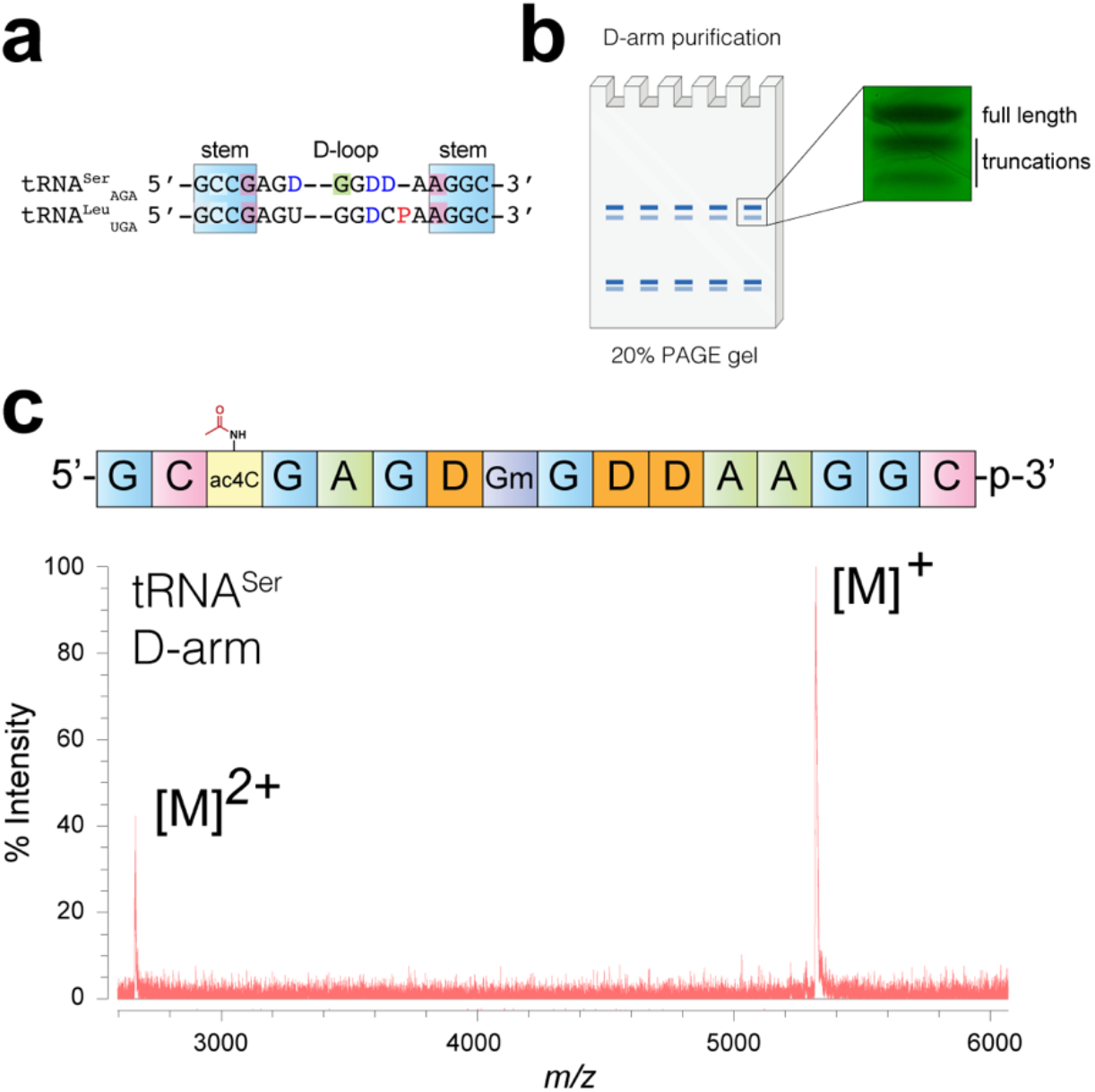
(a) Sequence alignment of D-arm of serine and leucine tRNAs. D = dihydrouridine, yellow = ac4C, green = Gm, red = purine-purine pair. See Table 2 for secondary structure. (c) PAGE analysis of crude product. (c) MALDI-TOF analysis of purified tRNA^Ser^ D-arm.

**Figure 6.**
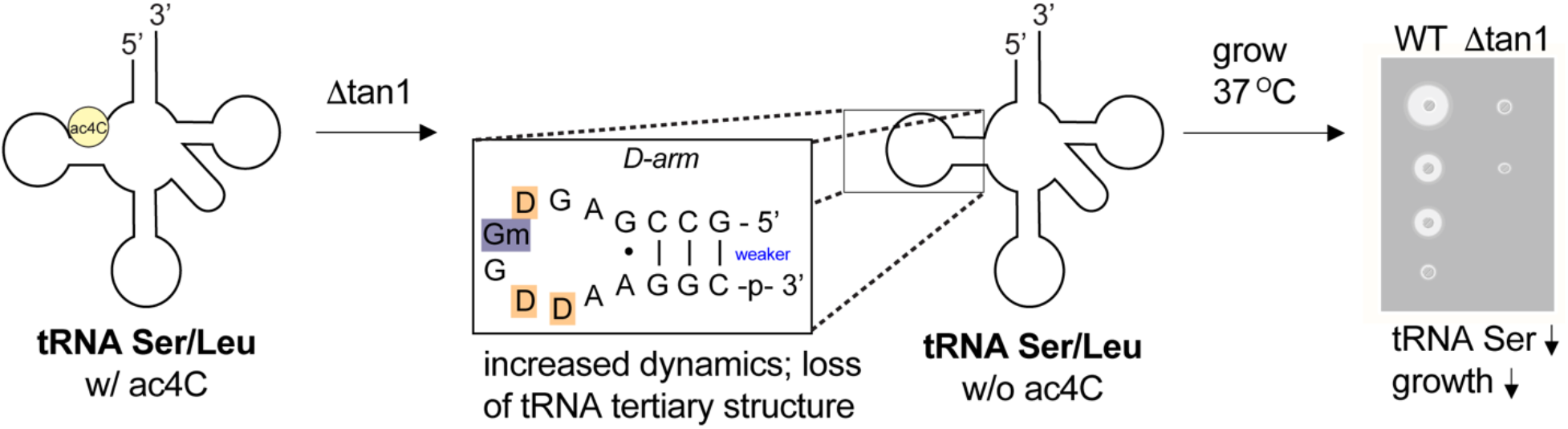
Model for regulation of tRNA stability by ac4C-dependent D-stem stabilization. Primary data demonstrating reduced levels of tRNA^Ser^ and reduced cell fitness in ac4C-deficient Δtan1 strains are provided in reference 35.

Human and yeast tRNA^Ser^_CGA_ share an identical sequence and modification profile in their D-arm, which is composed of a 4-bp stem that contains an internal ac4C-G and an 8-bp D-loop with three dihydrouridines (D) and one 2’-O-methylguanosine (Gm).^1,3,37^ Its synthesis presents a challenge due to its length (16 bp) and the presence of an additional labile nucleobase, D, that is prone to ring-opening. Previous studies have shown D-containing RNA can be obtained using phenoxyacyl-protected nucleobases.^38^ This led us to hypothesize that the even gentler *N*-ceoc protecting group strategy would be compatible with D, while also facilitating incorporation of ac4C (and Gm) into the hypermodified tRNA^Ser^ hairpin. To obtain the necessary building blocks, 5’-O-DMT-protected phosphoramidite monomers of D and Gm were synthesized. The synthesis of protected D used an adaptation of a previously reported method (Figure S6),^38^ while the Gm monomer was readily obtained via nucleobase deprotection of commercial starting material in a single step.^39^ These materials were then applied in combination with the previously described building blocks using the optimized solid-phase protocol to synthesize tRNA^Ser^ D-arm models containing either C or ac4C at the C12 position.

**Table 2.** UV-melting of synthetic RNA hairpin corresponding to the D-arm of human tRNA^Ser^. Left: schematic of tRNA^Ser^, showing site of ac4C at C12 in the stem of the D-loop. Right: Curves fit to melting data of C and ac4C-containing human tRNA^Ser^ D-arm hairpin. Bottom: Thermodynamic parameters obtained from UV melting experiments (n = 3). Full melting curves are provided in the Supplementary Information.

| variable | T<sub>m</sub> (°C) [5 μM] | ΔT<sub>m</sub> (°C) (ac4C) | ΔG (kcal/mol) | ΔH (kcal/mol) | ΔS (kcal/mol) |
| --- | --- | --- | --- | --- | --- |
| C | 62.9 |  | -2.4 ± 0.1 | -86 ± 3 | -29 ± 1 |
|  |  | +8.2 |  |  |  |
| ac4C | 71.1 |  | -3.4 ± 0.2 | -89 ± 7 | -32 ± 3 |

Analysis of crude reaction products revealed higher amounts of truncation products during the synthesis of tRNA hairpins relative to our rRNA-derived 10-mer, consistent with its longer linear sequence (Figure 5b). PAGE purification, and in the case of ac4C subsequent HPLC-purification, yielded the desired C- and ac4C-containing tRNA^Ser^ D-arm hairpins in quantities sufficient for biophysical characterization (Figure 5c).

Melting temperature measurements were performed at higher concentrations to account for the short stem structure and presence of non-aromatic nucleobases in the tRNA hairpin. Thermal denaturation analysis revealed a clear helix to coil transition for the ac4C-containing RNA at 71.4 ± 0.4 °C, while the non-acetylated hairpin melted at 62.9 ± 0.7 °C (ΔTm_ac4C v. C_ = +8.2 °C) (Table 2). This represents a stabilization of ~1 kcal/mol, similar in magnitude to the free energy change caused by inserting pseudouridine into a base-paired duplex.^40^ The UV melting profile of tRNA^Ser^ was not sensitive to concentration, consistent with a unimolecular (hairpin) as opposed to bimolecular (duplex) process. Across evolution, serine and leucine tRNAs are characterized by two unique elements: a variable region of more than 10 nucleotides and a conserved purine-purine (G13●A23) pair.^41^ Of note, these features converge on the tertiary structure of the tRNA formed by the D-arm. Our studies suggest ac4C may constitute a third distinct functional element in eukaryotic tRNA^Ser^ and tRNA^Leu^, and support the plausibility of a mechanism whereby cytidine acetylation regulates tRNA half-life and overall fitness by modulating the structural dynamics of these non-coding RNAs at elevated temperatures.

## Conclusion

Recent evidence linking Nat10 to disease has invigorated the study of cytidine acetylation in RNA. However, the precise effects of ac4C on nucleic acid structure and function remain unknown.*^17^* Here we report the synthesis and evaluation of *N*^4^-acetylcytidine in its physiological RNA sequence context. Systematic development of a mild non-nucleophilic RNA synthesis enabled the preparation of homogenous ac4C oligonucleotides. In a duplex RNA based on helix 45 of human 18S rRNA, we find that ac4C increases C-G base pair stability, an effect that is slightly augmented by the presence of a physiological G●U pair proximal to the acetylated 5’-C**C**G-3. Our synthetic method also provides access to the hypermodified D-arm hairpin of eukaryotic tRNA^Ser^ and we find that ac4C is highly stabilizing in this context (ΔTm_ac4C v. C_ = +8.2 °C). Previous studies have shown that destabilization of the D-stem can propagate in a zipper-like manner toward the anticodon arm,^42^ triggering disruption of tRNA tertiary structure and recognition by decay machinery.^43^ By providing empirical evidence that D-arm stabilization is highly dependent on ac4C at elevated temperatures, our studies differentiate the catalytic and non-catalytic functions of the Kre33/Tan1 complex, and provide a molecular rationale for why Δtan1 strains exhibit decreased tRNA^Ser^ and reduced fitness under environmental stress.

Clarifying the thermodynamic consequences of ac4C in these physiologically-relevant sequence contexts also raises new questions. First, how does ac4C stabilize duplex RNA? Comparative cryo-EM analyses of an archaeal ribosome with >100 sites of ac4C did not observe large perturbations of hydrogen-bonding when this modification was deleted (Figure S7), albeit at limited resolution. One source of stability may come from the exocyclic acetyl group of ac4C, which projects into the major groove of duplex RNA and has been hypothesized to contribute to binding enthalpy by serving as a stable covalent replacement for ordered waters at elevated temperatures.^6^ Another analogy may be found in 5-formylcytidine, which presents an exocyclic hydrogen bonding network towards the C-H edge that is similar to the favored conformation of ac4C, and has been shown to increase base-stacking.^44^ Understanding these interactions will benefit from additional biophysical interrogation and higher resolution structural analyses, both of which will be facilitated by our method.

A second question is: why is cytidine acetylation restricted to tRNA^Ser^ and tRNA^Leu^? As noted above, these species are distinguished from other tRNAs by large variable regions (>10 nt) and the presence of a purine-purine (G●A) pair in the D-stem.^41^ Here we suggest two hypotheses. These tRNAs could be uniquely recognized as substrates by the Nat10/Thumpd1 (Kre33/Tan1) complex. Alternatively, these tRNAs could be uniquely susceptible to modification-dependent structural stabilization, given the rare occurrence of a non-canonical G●A pair directly adjacent to ac4C.^45,46^ Previous studies have demonstrated the G●A interaction is highly sensitive to sequence context.^47^ Understanding how RNA modifications affect adjacent non-canonical base pairs is an important question posed by our observations, which future research will seek to address.

Finally, we anticipate the synthetic method described here will enable several additional exciting applications. Besides the D-arm of eukaryotic tRNA^Ser/Leu^, hypermodified ac4C-containing RNAs are also present in the bacterial anticodon arm and the archaeal ribosome (Figure S8).^31,48^ The latter also contains 2’-O-methylated ac4C (ac4Cm), the so-called “the most conformationally rigid nucleobase” present in nature.^49^ Our methods should facilitate pioneering biophysical and structural analyses of these modification-rich contexts. In addition, the ability to site-specifically incorporate ac4C opens the door to exploring its effects in functional nucleic acids, including short guide RNAs, short interfering RNAs, and antisense oligonucleotides. A recent study found homogenous replacement of cytidine with ac4C reduced the immunogenicity of synthetic mRNAs,^18^ suggesting this modification’s potential therapeutic utility. We envision our synthetic route as being amenable to incorporation of diverse *N*^4^-acylated cytidines^50^ as well as other electrophilic nucleobases,^51^ further extending the chemical functionalities that may be explored in these applications. By illuminating how ac4C influences duplex RNA stability in physiological sequence contexts, this chemistry provides a foundation for understanding and exploiting cytidine acetylation as a novel regulatory element in biology, biotechnology, and disease.

## Supporting Information

Supporting Information including supplemental figures and tables and materials and methods are available free of charge on the ACS publications website at pubs.acs.org.

## Acknowledgements

The authors thank Prof. Moran Shalev-Benami (Weizmann Institute) and Prof. Marc Greenberg (Johns Hopkins University) for helpful discussions. We thank the Biophysics Resource in the Center for Structural Biology, Center for Cancer Research, NCI at Frederick for assistance with high resolution LC-MS characterization and UV melting studies. Figures created with help of Biorender.com. This work was supported by the Intramural Research Program of the NIH, National Cancer Institute, Center for Cancer Research (ZIA-BC011488-05).

## References

(1) Boccaletto, P.; Machnicka, M. A.; Purta, E.; Piątkowski, P.; Bagiński, B.; Wirecki, T. K.; de Crécy-Lagard, V.; Ross, R.; Limbach, P. A.; Kotter, A.; Helm, M.; Bujnicki, J. M. MODOMICS: A Database of RNA Modification Pathways. 2017 Update. Nucleic Acids Research. 2018, pp D303–D307. https://doi.org/10.1093/nar/gkx1030.

(2) Zachau, H. G.; Dütting, D.; Feldmann, H. Nucleotide Sequences of Two Serine-Specific Transfer Ribonucleic Acids. Angew. Chem. Int. Ed Engl. 1966, 5 (4), 422.

(3) Staehelin, M.; Rogg, H.; Baguley, B. C.; Ginsberg, T.; Wehrli, W. Structure of a Mammalian Serine tRNA. Nature. 1968, pp 1363–1365. https://doi.org/10.1038/2191363a0.

(4) Ito, S.; Horikawa, S.; Suzuki, T.; Kawauchi, H.; Tanaka, Y.; Suzuki, T.; Suzuki, T. Human NAT10 Is an ATP-Dependent RNA Acetyltransferase Responsible for N4-Acetylcytidine Formation in 18 S Ribosomal RNA (rRNA). J. Biol. Chem. 2014, 289 (52), 35724–35730.

(5) Sharma, S.; Langhendries, J.-L.; Watzinger, P.; Kötter, P.; Entian, K.-D.; Lafontaine, D. L. J. Yeast Kre33 and Human NAT10 Are Conserved 18S rRNA Cytosine Acetyltransferases That Modify tRNAs Assisted by the Adaptor Tan1/THUMPD1. Nucleic Acids Res. 2015, 43 (4), 2242–2258.

(6) Sas-Chen, A.; Thomas, J. M.; Matzov, D.; Taoka, M.; Nance, K. D.; Nir, R.; Bryson, K. M.; Shachar, R.; Liman, G. L. S.; Burkhart, B. W.; Gamage, S. T.; Nobe, Y.; Briney, C. A.; Levy, M. J.; Fuchs, R. T.; Robb, G. B.; Hartmann, J.; Sharma, S.; Lin, Q.; Florens, L.; Washburn, M. P.; Isobe, T.; Santangelo, T. J.; Shalev-Benami, M.; Meier, J. L.; Schwartz, S. Dynamic RNA Acetylation Revealed by Quantitative Cross-Evolutionary Mapping. Nature 2020, 583 (7817), 638–643.

(7) Larrieu, D.; Britton, S.; Demir, M.; Rodriguez, R.; Jackson, S. P. Chemical Inhibition of NAT10 Corrects Defects of Laminopathic Cells. Science 2014, 344 (6183), 527–532.

(8) Tschida, B. R.; Temiz, N. A.; Kuka, T. P.; Lee, L. A.; Riordan, J. D.; Tierrablanca, C. A.; Hullsiek, R.; Wagner, S.; Hudson, W. A.; Linden, M. A.; Amin, K.; Beckmann, P. J.; Heuer, R. A.; Sarver, A. L.; Yang, J. D.; Roberts, L. R.; Nadeau, J. H.; Dupuy, A. J.; Keng, V. W.; Largaespada, D. A. Insertional Mutagenesis in Mice Identifies Drivers of Steatosis-Associated Hepatic Tumors. Cancer Res. 2017, 77 (23), 6576–6588.

(9) Parthasarathy, R.; Ginell, S. L.; De, N. C.; Chheda, G. B. Conformation of N4-Acetylcytidine, a Modified Nucleoside of tRNA, and Stereochemistry of Codon-Anticodon Interaction. Biochem. Biophys. Res. Commun. 1978, 83 (2), 657–663.

(10) Kumbhar, B. V.; Kamble, A. D.; Sonawane, K. D. Conformational Preferences of Modified Nucleoside N(4)-Acetylcytidine, ac4C Occur at “Wobble” 34th Position in the Anticodon Loop of tRNA. Cell Biochem. Biophys. 2013, 66 (3), 797–816.

(11) Harcourt, E. M.; Kietrys, A. M.; Kool, E. T. Chemical and Structural Effects of Base Modifications in Messenger RNA. Nature 2017, 541 (7637), 339–346.

(12) Natchiar, S. K.; Myasnikov, A. G.; Kratzat, H.; Hazemann, I.; Klaholz, B. P. Visualization of Chemical Modifications in the Human 80S Ribosome Structure. Nature 2017, 551 (7681), 472–477.

(13) Coureux, P.-D.; Lazennec-Schurdevin, C.; Bourcier, S.; Mechulam, Y.; Schmitt, E. Cryo-EM Study of an Archaeal 30S Initiation Complex Gives Insights into Evolution of Translation Initiation. Commun Biol 2020, 3 (1), 58.

(14) Thalalla Gamage, S.; Sas-Chen, A.; Schwartz, S.; Meier, J. L. Quantitative Nucleotide Resolution Profiling of RNA Cytidine Acetylation by ac4C-Seq. Nat. Protoc. 2021, 16 (4), 2286–2307.

(15) Orita, I.; Futatsuishi, R.; Adachi, K.; Ohira, T.; Kaneko, A.; Minowa, K.; Suzuki, M.; Tamura, T.; Nakamura, S.; Imanaka, T.; Suzuki, T.; Fukui, T. Random Mutagenesis of a Hyperthermophilic Archaeon Identified tRNA Modifications Associated with Cellular Hyperthermotolerance. Nucleic Acids Res. 2019, 47 (4), 1964–1976.

(16) Kowalak, J. A.; Dalluge, J. J.; McCloskey, J. A.; Stetter, K. O. The Role of Posttranscriptional Modification in Stabilization of Transfer RNA from Hyperthermophiles. Biochemistry 1994, 33 (25), 7869–7876.

(17) Sinclair, W. R.; Arango, D.; Shrimp, J. H.; Zengeya, T. T.; Thomas, J. M.; Montgomery, D. C.; Fox, S. D.; Andresson, T.; Oberdoerffer, S.; Meier, J. L. Profiling Cytidine Acetylation with Specific Affinity and Reactivity. ACS Chem. Biol. 2017, 12 (12), 2922–2926.

(18) Nance, K. D.; Gamage, S. T.; Alam, M. M.; Yang, A.; Levy, M. J.; Link, C. N.; Florens, L.; Washburn, M. P.; Gu, S.; Oppenheim, J. J.; Meier, J. L. Cytidine Acetylation Yields a Hypoinflammatory Synthetic Messenger RNA. Cell Chemical Biology. 2021. https://doi.org/10.1016/j.chembiol.2021.07.003.

(19) Roy, S.; Caruthers, M. Synthesis of DNA/RNA and Their Analogs via Phosphoramidite and H-Phosphonate Chemistries. Molecules 2013, 18 (11), 14268–14284.

(20) Schulhof, J. C.; Molko, D.; Teoule, R. The Final Deprotection Step in Oligonucleotide Synthesis Is Reduced to a Mild and Rapid Ammonia Treatment by Using Labile Base-Protecting Groups. Nucleic Acids Res. 1987, 15 (2), 397–416.

(21) Hu, B.; Zhong, L.; Weng, Y.; Peng, L.; Huang, Y.; Zhao, Y.; Liang, X.-J. Therapeutic siRNA: State of the Art. Signal Transduction and Targeted Therapy. 2020. https://doi.org/10.1038/s41392-020-0207-x.

(22) Xu, J.; Duffy, C. D.; Chan, C. K. W.; Sutherland, J. D. Solid-Phase Synthesis and Hybrization Behavior of Partially 2′/3′-O-Acetylated RNA Oligonucleotides. The Journal of Organic Chemistry. 2014, pp 3311–3326. https://doi.org/10.1021/jo5002824.

(23) Bowler, F. R.; Chan, C. K. W.; Duffy, C. D.; Gerland, B.; Islam, S.; Powner, M. W.; Sutherland, J. D.; Xu, J. Prebiotically Plausible Oligoribonucleotide Ligation Facilitated by Chemoselective Acetylation. Nat. Chem. 2013, 5 (5), 383–389.

(24) Merk, C.; Reiner, T.; Kvasyuk, E.; Pfleiderer, W. Nucleotides, Part LXVII, The 2-Cyanoethyl and (2-Cyanoethoxy)carbonyl Group for Base Protection in Nucleoside and Nucleotide Chemistry. Helvetica Chimica Acta. 2000, pp 3198–3210. https://doi.org/10.1002/1522-2675(20001220)83:12<3198::aid-hlca3198>3.0.co;2-q.

(25) Serebryany, V.; Beigelman, L. An Efficient Preparation of Protected Ribonucleosides for Phosphoramidite RNA Synthesis. Tetrahedron Letters. 2002, pp 1983–1985. https://doi.org/10.1016/s0040-4039(02)00181-8.

(26) Miller, N.; Cerutti, P. The Synthesis of N4-Acetyl-3,4,5,6-Tetrahydrocytidine and Copolymers of Cytidylic Acid and N4-Acetyl-3,4,5,6-Tetrahydrocytidylic Acid. Journal of the American Chemical Society. 1967, pp 2767–2768. https://doi.org/10.1021/ja00987a065.

(27) Helene, C.; Douzou, P.; Michelson, A. M. Energy Transfer in Dinucleotides. Proceedings of the National Academy of Sciences. 1966, pp 376–381. https://doi.org/10.1073/pnas.55.2.376.

(28) Zhu, Q.; Delaney, M. O.; Greenberg, M. M. Observation and Elimination of N-Acetylation of Oligonucleotides Prepared Using Fast-Deprotecting Phosphoramidites and Ultra-Mild Deprotection. Bioorg. Med. Chem. Lett. 2001, 11 (9), 1105–1107.

(29) Ohkubo, A.; Sakamoto, K.; Miyata, K.-I.; Taguchi, H.; Seio, K.; Sekine, M. Convenient Synthesis of N-Unprotected Deoxynucleoside 3’-Phosphoramidite Building Blocks by Selective Deacylation of N-Acylated Species and Their Facile Conversion to Other N-Functionalized Derivatives. Org. Lett. 2005, 7 (24), 5389–5392.

(30) Wada, T.; Kobori, A.; Kawahara, S.-I.; Sekine, M. Synthesis and Properties of Oligodeoxyribonucleotides Containing 4-N-Acetylcytosine Bases. Tetrahedron Letters. 1998, pp 6907–6910. https://doi.org/10.1016/s0040-4039(98)01449-x.

(31) Stern, L.; Schulman, L. H. The Role of the Minor Base N4-Acetylcytidine in the Function of the Escherichia Coli Noninitiator Methionine Transfer RNA. J. Biol. Chem. 1978, 253 (17), 6132–6139.

(32) Oashi, Z.; Murao, K.; Yahagi, T.; Von Minden, D. L.; McCloskey, J. A.; Nishimura, S. Characterization of C + Located in the First Position of the Anticodon of Escherichia Coli tRNA Met as N 4 −Acetylcytidine. Biochim. Biophys. Acta 1972, 262 (2), 209–213.

(33) Varani, G.; McClain, W. H. The G·U Wobble Base Pair. EMBO reports. 2000, pp 18–23. https://doi.org/10.1093/embo-reports/kvd001.

(34) Johansson, M. J. O. The Saccharomyces Cerevisiae TAN1 Gene Is Required for N4-Acetylcytidine Formation in tRNA. RNA. 2004, pp 712–719. https://doi.org/10.1261/rna.5198204.

(35) Dewe, J. M.; Whipple, J. M.; Chernyakov, I.; Jaramillo, L. N.; Phizicky, E. M. The Yeast Rapid tRNA Decay Pathway Competes with Elongation Factor 1A for Substrate tRNAs and Acts on tRNAs Lacking One or More of Several Modifications. RNA 2012, 18 (10), 1886–1896.

(36) Roovers, M. A Primordial RNA Modification Enzyme: The Case of tRNA (m1A) Methyltransferase. Nucleic Acids Research. 2004, pp 465–476. https://doi.org/10.1093/nar/gkh191.

(37) Jühling, F.; Mörl, M.; Hartmann, R. K.; Sprinzl, M.; Stadler, P. F.; Pütz, J. tRNAdb 2009: Compilation of tRNA Sequences and tRNA Genes. Nucleic Acids Res. 2009, 37 (Database issue), D159–D162.

(38) Dyubankova, N.; Sochacka, E.; Kraszewska, K.; Nawrot, B.; Herdewijn, P.; Lescrinier, E. Contribution of Dihydrouridine in Folding of the D-Arm in tRNA. Org. Biomol. Chem. 2015, 13 (17), 4960–4966.

(39) Ohkubo, A.; Kuwayama, Y.; Kudo, T.; Tsunoda, H.; Seio, K.; Sekine, M. O-Selective Condensation Using P-N Bond Cleavage in RNA Synthesis without Base Protection. Org. Lett. 2008, 10 (13), 2793–2796.

(40) Meroueh, M.; Grohar, P. J.; Qiu, J.; SantaLucia, J., Jr; Scaringe, S. A.; Chow, C. S. Unique Structural and Stabilizing Roles for the Individual Pseudouridine Residues in the 1920 Region of Escherichia Coli 23S rRNA. Nucleic Acids Res. 2000, 28 (10), 2075–2083.

(41) Giegé, R.; Puglisi, J. D.; Florentz, C. tRNA Structure and Aminoacylation Efficiency. Prog. Nucleic Acid Res. Mol. Biol. 1993, 45, 129–206.

(42) Bonnefond, L.; Florentz, C.; Giegé, R.; Rudinger-Thirion, J. Decreased Aminoacylation in Pathology-Related Mutants of Mitochondrial tRNATyr Is Associated with Structural Perturbations in tRNA Architecture. RNA 2008, 14 (4), 641–648.

(43) Guy, M. P.; Young, D. L.; Payea, M. J.; Zhang, X.; Kon, Y.; Dean, K. M.; Grayhack, E. J.; Mathews, D. H.; Fields, S.; Phizicky, E. M. Identification of the Determinants of tRNA Function and Susceptibility to Rapid tRNA Decay by High-Throughput in Vivo Analysis. Genes Dev. 2014, 28 (15), 1721–1732.

(44) Wang, R.; Luo, Z.; He, K.; Delaney, M. O.; Chen, D.; Sheng, J. Base Pairing and Structural Insights into the 5-Formylcytosine in RNA Duplex. Nucleic Acids Res. 2016, 44 (10), 4968–4977.

(45) Dock-Bregeon, A. C.; Westhof, E.; Giegé, R.; Moras, D. Solution Structure of a tRNA with a Large Variable Region: Yeast tRNASer. J. Mol. Biol. 1989, 206 (4), 707–722.

(46) Biou, S.; Cusack, V.; Yaremchuk, A.; Tukalo, M. THE 2.9 ANGSTROMS CRYSTAL STRUCTURE OF T. THERMOPHILUS SERYL-TRNA SYNTHETASE COMPLEXED WITH TRNA SER. 1994. https://doi.org/10.2210/pdb1ser/pdb.

(47) Schroeder, S.; Kim, J.; Turner, D. H. G.A and U.U Mismatches Can Stabilize RNA Internal Loops of Three Nucleotides. Biochemistry 1996, 35 (50), 16105–16109.

(48) Taniguchi, T.; Miyauchi, K.; Sakaguchi, Y.; Yamashita, S.; Soma, A.; Tomita, K.; Suzuki, T. Acetate-Dependent tRNA Acetylation Required for Decoding Fidelity in Protein Synthesis. Nat. Chem. Biol. 2018, 14 (11), 1010–1020.

(49) Bruenger, E.; Kowalak, J. A.; Kuchino, Y.; McCloskey, J. A.; Mizushima, H.; Stetter, K. O.; Crain, P. F. 5S rRNA Modification in the Hyperthermophilic Archaea Sulfolobus Solfataricus and Pyrodictium Occultum. FASEB J. 1993, 7 (1), 196–200.

(50) Wada, T.; Kobori, A.; Kawahara, S.-I.; Sekine, M. Synthesis and Hybridization Ability of Oligodeoxyribonucleotides Incorporating N-Acyldeoxycytidine Derivatives. European Journal of Organic Chemistry. 2001, p 4583. https://doi.org/10.1002/1099-0690(200112)2001:24<4583::aid-ejoc4583>3.0.co;2-r.

(51) Dai, W.; Li, A.; Yu, N. J.; Nguyen, T.; Leach, R. W.; Wühr, M.; Kleiner, R. E. Activity-Based RNA-Modifying Enzyme Probing Reveals DUS3L-Mediated Dihydrouridylation. Nat. Chem. Biol. 2021. https://doi.org/10.1038/s41589-021-00874-8.

