## Supplementary Information for "Site-Specific Synthesis of N4-Acetylcytidine in RNA Reveals Physiological Duplex Stabilization"

**Table of Contents for Supporting Information**


| | Page |
|---|---|
| Figure S1: DBU treatment of ac4C in the presence of morpholine | S2 |
| Figure S2: Analysis of pivalic anhydride capping in crude RNA cleavages | S3 |
| Figure S3: Optimization of RNA deprotection | S4 |
| Figure S4: PAGE purification and MALDI of rRNA decamer | S5 |
| Figure S5: C•U wobble pair in *T. kodakarensis* rRNA | S6 |
| Figure S6: Synthesis of dihydrouridine (D) phosphoramidite | S7 |
| Figure S7: Comparison of *T. kodakarensis* rRNA ± ac4C | S8 |
| Figure S8: Exemplary hypermodified ac4C sequence contexts | S9 |
| Synthetic procedures | S10-25 |
| UV melting analysis | S26 |
| UV melting curves and van't Hoff plots | S26-30 |
| References | S31 |

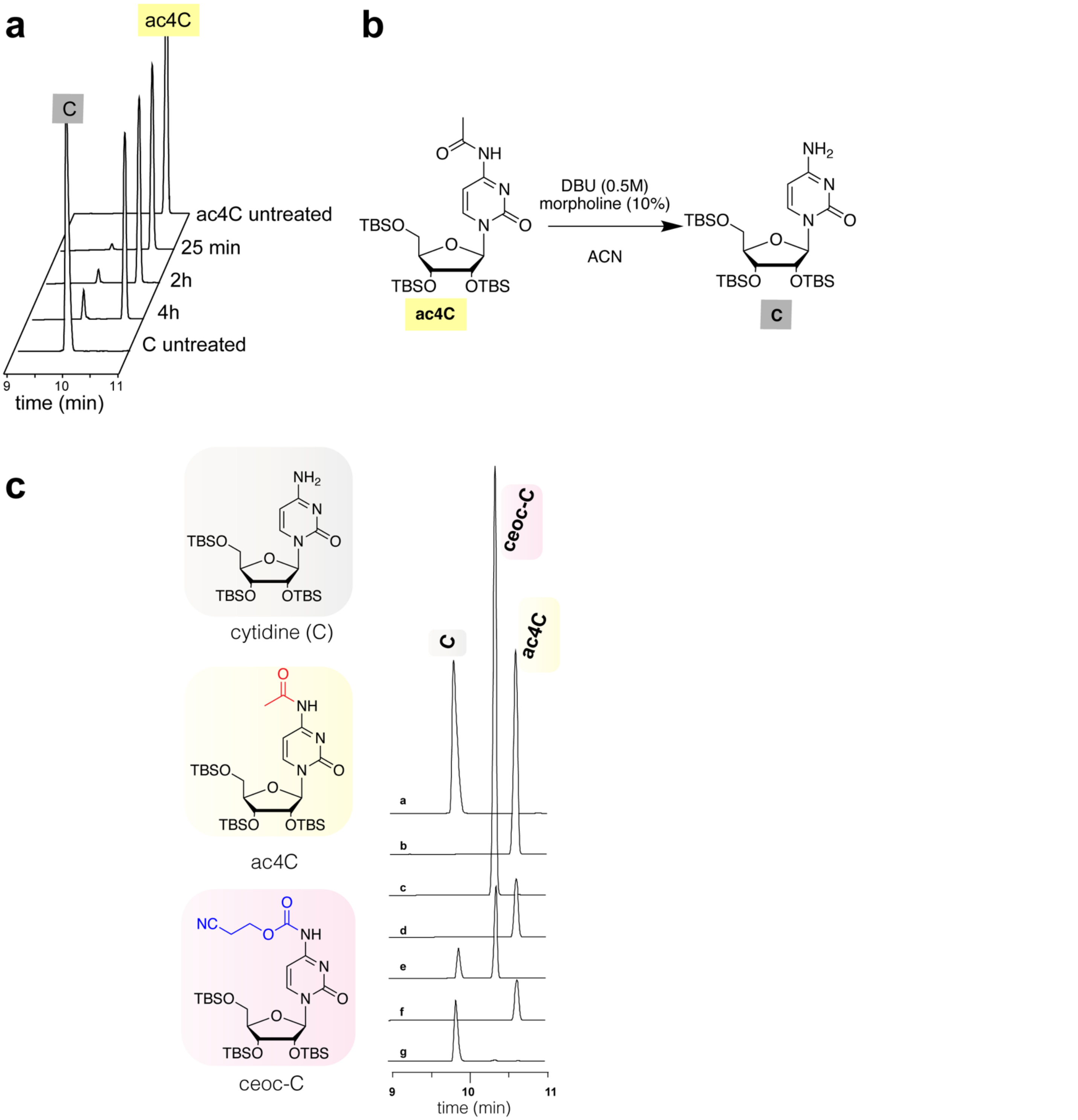


**Figure S1**. (a-b) DBU treatment of model ac4C compound with 10% morpholine. Previous studies of electrophilic RNAs (2'-O-acetyl esters) have used morpholine as a nucleophilic scavenger to prevent alkylation byproducts caused by acrylonitrile release.[1] Incompatibility with ac4C highlights the augmented sensitivity of this modification. (c) Extended HPLC traces from Figure 2. Conditions: a: C standard; b: ac4C standard; c: ceoc-C standard; d: ac4C after 25 min of DBU treatment; e: ceoc-C after 25 min of DBU treatment. F: ac4C after 4 h of DBU treatment; g: ceoc-C after 4 h of DBU treatment.

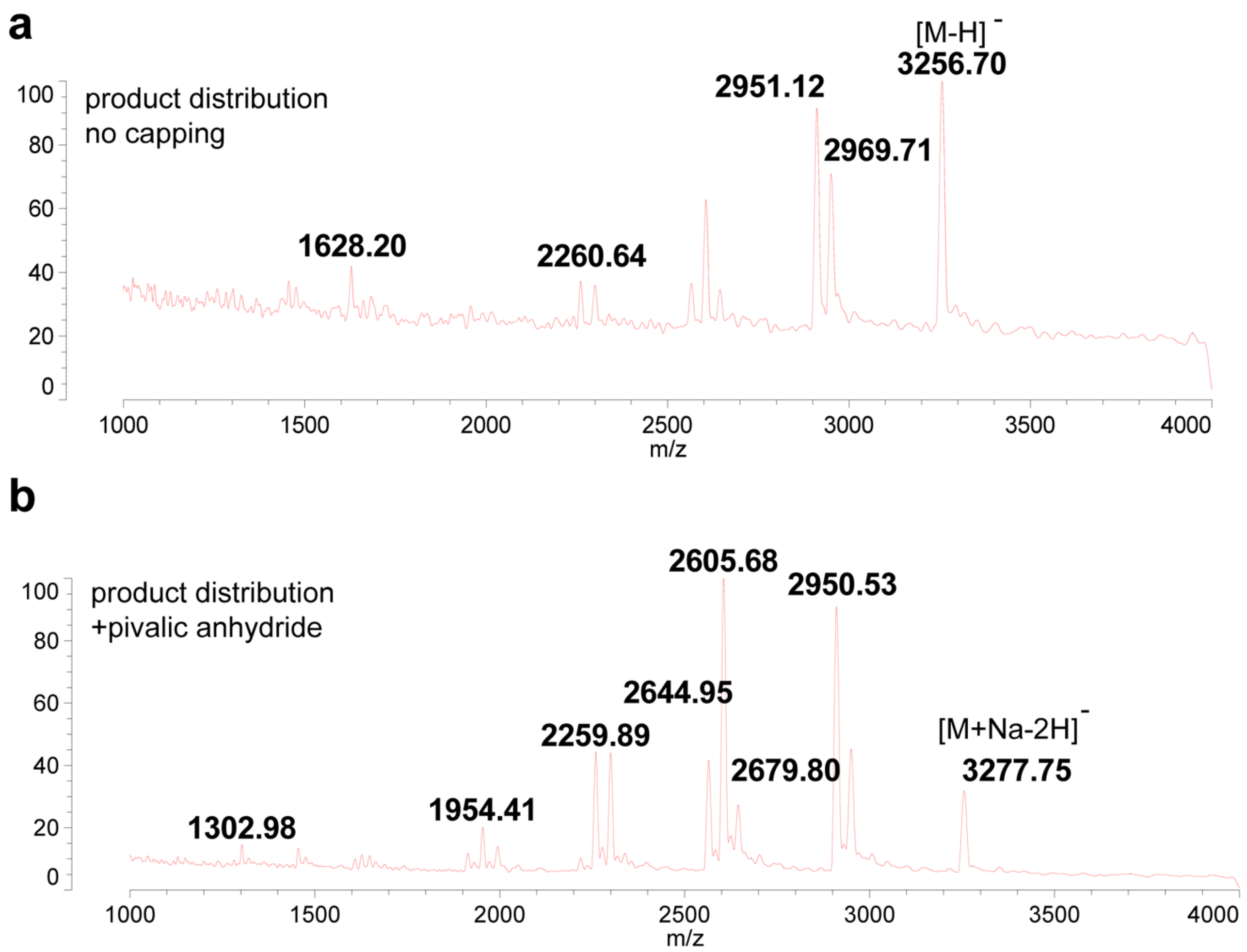


**Figure S2**. MALDI-TOF analysis of product distribution in crude RNA synthesis products. Crude products were cleaved from resin and deprotected but not subjected to further purification. (a) Product distribution without 5'-OH capping step; (b) Product distribution using pivalic anhydride as a 5'-OH capping agent. Full length product (5'-CUUC(ac4C)GUAGGp-3') m/z calc'd as $[M-H]^-$ of 3256.4, $[M+Na-2H]^-$ of 3278.4.

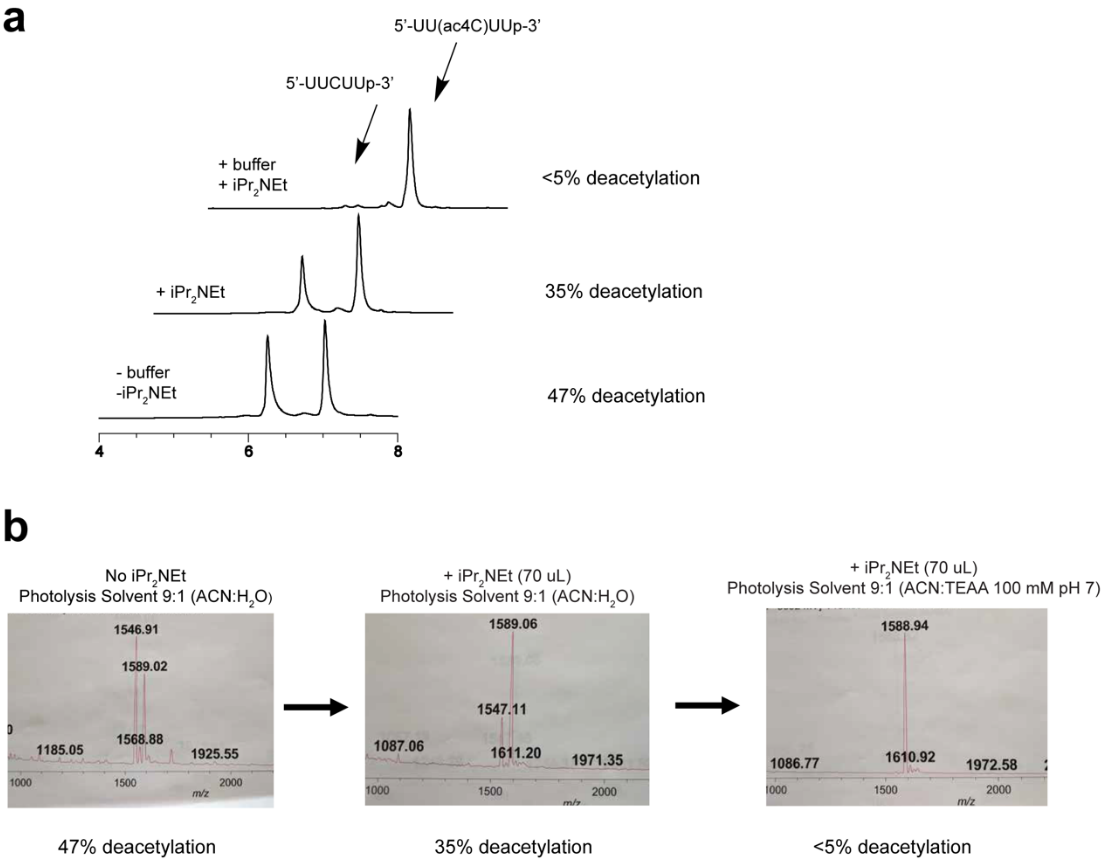


**Figure S3**. Optimization of RNA deprotection.. (a) HPLC chromatograms of crude ac4C RNA oligonucleotides with no modifications to previously reported photolysis and 2'-OH silyl deprotections steps (top), following addition of $iPr_2NEt$ to 2'-OH silyl deprotections step (middle), and following addition of $iPr_2NEt$ to 2'-OH silyl deprotections step and triethylammonium acetate buffer to the photolysis step (bottom). (b) MALDI spectra of HPLC-purified samples prepared in panel a indicates the improved purity by reducting byproduct formation.

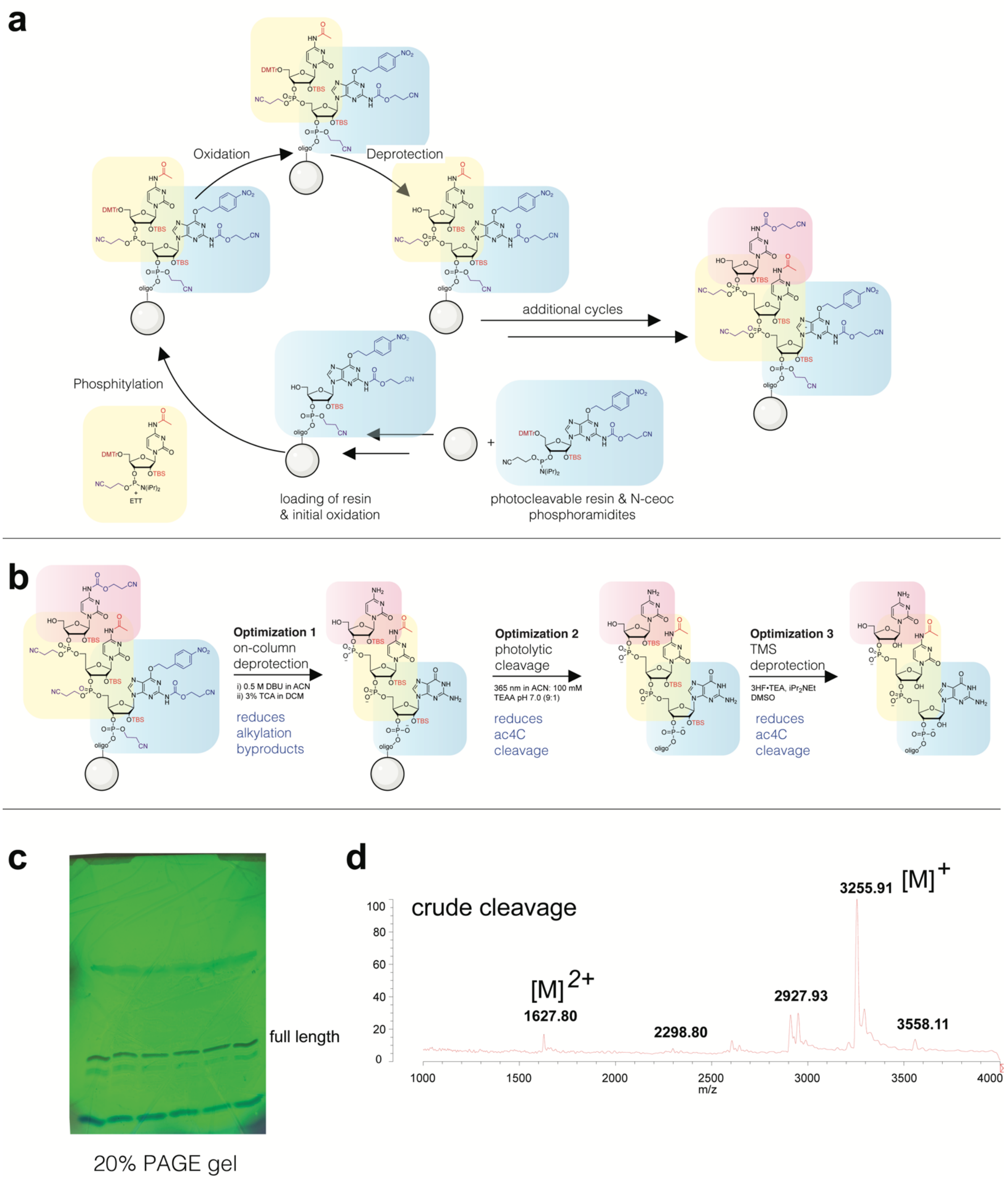


**Figure S4**. PAGE purification and MALDI ac4C decamer. (a-b) Chemical structures corresponding to RNA synthesis abbreviated in Figure 4. (c) Polyacrylamide gel (20%) purification of crude ac4C-containing RNA oligonucleotide (5'-CUUC(ac4C)GUAGGp-3'). (d) MALDI spectrum of crude RNA prior to PAGE purification.

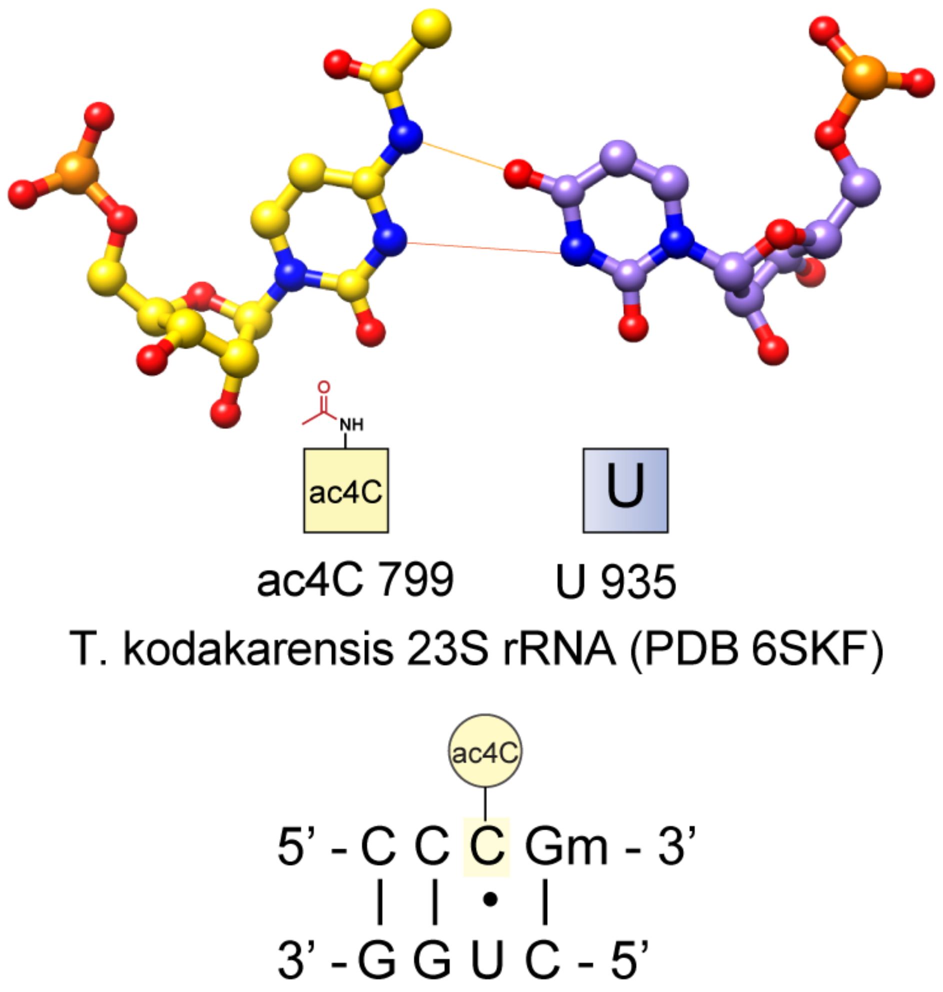


**Figure S5**. An ac4C•U wobble pair observed in *T. kodakarensis* rRNA (PDB 6SKF).[2]

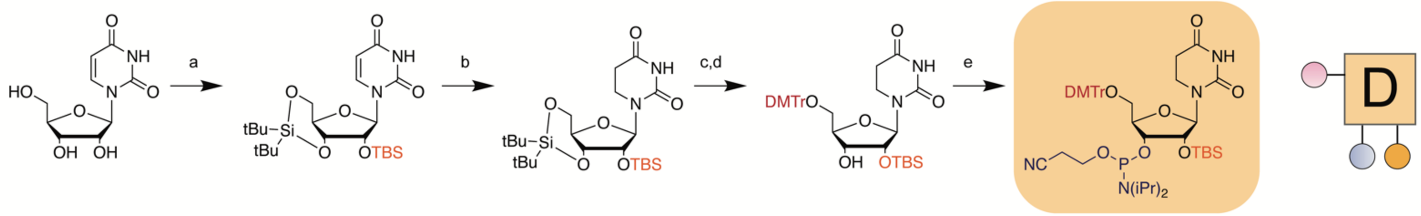


**Figure S6**. Synthesis of dihydrouridine phosphoramidite. (a) i) $(O\text{-}tBu)_2Si(OTf)_2$, DMF, 0 °C, ii) TBS-Cl, imidazole, DMF, 60 °C; (b) $H_2$ (1 atm), Rh/alumina, MeOH, 23 °C; (c) HF-pyridine, pyridine, DCM, 0 °C; (d) DMTr-Cl, pyridine, 4 °C; (e) $(OCH_2CH_2CN)(iPr_2N)PCl$, $iPr_2NEt$, THF, 23 °C;

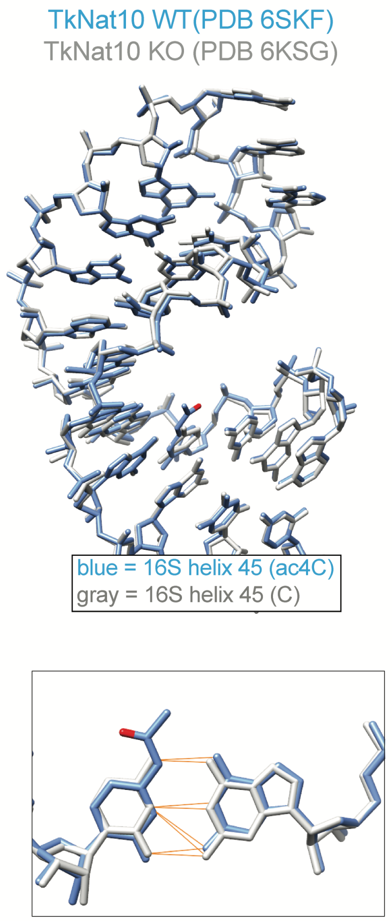


**Figure S7**. Top: Structural overlay of *T. kodakarensis* rRNA helix 45 rRNA with and without ac4C (PDBs: 6SKF, 6KSG Bottom: Zoomed-in overlay of ac4C-G and C-G base pairs in wild type and TkNat10 KO (ac4C-less) *T. kodakarensis* rRNA helix 45.

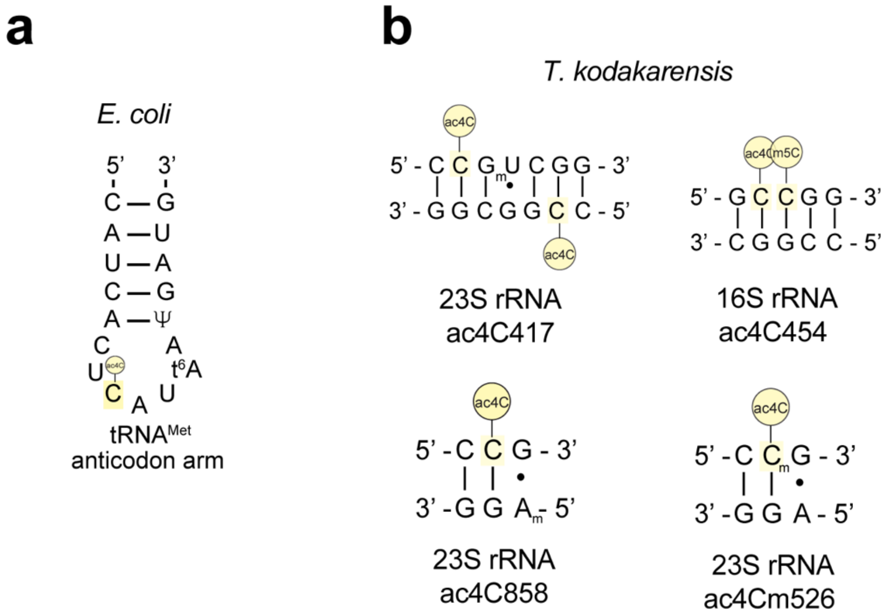


**Figure S8**. Examples of additional hypermodified RNA contexts that contain ac4C which are now accessible using the chemistry reported here. (a) The anticodon arm of *E. coli* tRNA$^{Met}$.[3] (b) Four sites in Thermococcus kodakarensis rRNA with multiple modifications including cytidine acetylation.

**Synthetic Procedures**

***General***

Unless otherwise noted, all reagents were obtained from commercial suppliers and used without further purification. Yields of all reactions refer to the purified products. Silica chromatography was carried out in the indicated solvent system using pre-packed silica gel cartridges for use on the Teledyne ISCO Purification System. $^{1}H$ and $^{13}C$ NMR spectra were acquired on a Bruker 500 MHz instrument operating at 500 MHz for $^{1}H$, 126 MHz for $^{13}C$, and 202 MHz for $^{31}P$. $^{31}P$ chemical shifts are reported relative to triphenylphosphine oxide at δ 24.7 as an external standard. NMR spectra were standardized to the NMR solvent signal as reported by Gottlieb.[4]

***Synthesis of models for deprotection study***

*N4-Acetylcytidine*

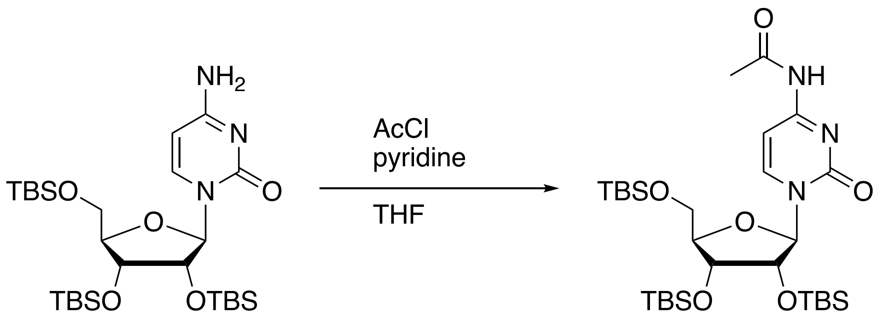


To a solution of protected cytidine (0.200 g, 0.34 mmol) in THF (3.4 mL) was added pyridine (0.06 mL, 0.68 mmol) followed by acetyl chloride (0.03 mL, 0.41 mmol). The reaction mixture was stirred at 23 °C for 16 h then diluted with diethyl ether (50 mL), transferred to a separatory funnel, and washed with water (3 x 50 mL). The organic layer was then dried over anhydrous $Na_2SO_4$, filtered, and then condensed under reduced pressure. Silica flash chromatography yielded the acetylated cytidine (0.158 g, 74% yield). $^{1}H$ NMR (500 MHz, $CDCl_3$) δ 10.43 (s, 1H), 8.57 (d, *J* = 7.5 Hz, 1H), 7.39 (d, *J* = 7.6 Hz, 1H), 5.76 (d, *J* = 1.3 Hz, 1H), 4.18 – 4.07 (m, 3H), 4.04 (dd, *J* = 7.9, 3.8 Hz, 1H), 3.79 (dd, *J* = 11.9, 1.5 Hz, 1H), 2.30 (s, 3H), 0.96 (s, 8H), 0.90 (s, 9H), 0.87 (s, 8H), 0.21 (s, 3H), 0.15 (s, 3H), 0.13 (s, 3H), 0.10 (s, 3H), 0.04 (s, 3H), 0.04 (s, 3H). $^{13}C$ NMR (126 MHz, $CDCl_3$) δ 171.70, 162.83, 154.46, 145.47, 96.23, 91.09, 83.19, 76.36, 68.92, 60.69, 26.22, 26.00, 25.98, 25.05, 18.69, 18.19, 18.17, -3.94, -4.03, -4.92, -5.05, -5.09, -5.42. HRMS (ESI/Q-TOF) calcd for $C_{29}H_{58}N_3O_6Si_3^+$ $[M + H]^+$ 628.3628, found 628.3629.

*Cytidine*

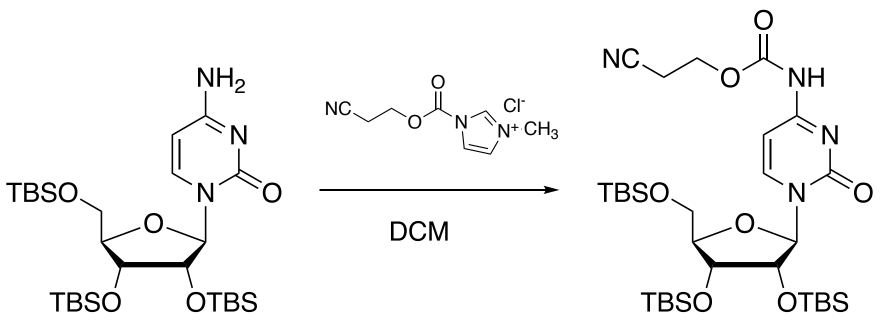


To a solution of protected cytidine (0.200 g, 0.34 mmol) in DCM (6.8 mL) was added ceoc-carbonyl-*N*-methylimidazolium chloride (0.093 g, 0.43 mmol). The reaction mixture was stirred at 23 °C for 16 h then diluted with DCM (50 mL), transferred to a separatory funnel, and washed with water (3 x 50 mL). The organic layer was then dried over anhydrous $Na_2SO_4$, filtered, and then condensed under reduced pressure. The resulting crude product was purified by silica flash chromatography to yield *N*-ceoc protected cytidine (0.145 g, 62% yield). $^1$H NMR (500 MHz, $CDCl_3$) δ 8.57 (s, 1H), 7.11 (s, 1H), 5.77 (s, 1H), 4.38 (qt, *J* = 11.0, 6.5 Hz, 2H), 4.17 – 4.11 (m, 2H), 4.14 – 4.07 (m, 1H), 4.04 (dd, *J* = 7.8, 3.8 Hz, 1H), 3.79 (dd, *J* = 11.9, 1.5 Hz, 1H), 2.78 (td, *J* = 6.5, 2.9 Hz, 2H), 0.96 (s, 9H), 0.91 (s, 9H), 0.88 (s, 9H), 0.23 (s, 3H), 0.15 (s, 3H), 0.13 (s, 3H), 0.12 (s, 3H), 0.08 (s, 0H), 0.04 (s, 6H). $^{13}$C NMR (126 MHz, $CDCl_3$) δ 162.21, 154.49, 152.04,145.32, 116.51, 91.01, 83.23, 76.31, 68.99, 60.75, 60.27, 26.21, 25.99, 25.98, 18.69, 18.19, 18.18, -3.93, -4.02, -4.91, -4.98, -5.10, -5.45. HRMS (ESI/Q-TOF) calcd for $C_{31}H_{59}N_4O_7Si_3^+$ [M + H]$^+$ 683.3686, found 683.3692.

### ***Synthesis of phosphoramidite building blocks***

*Adenosine*

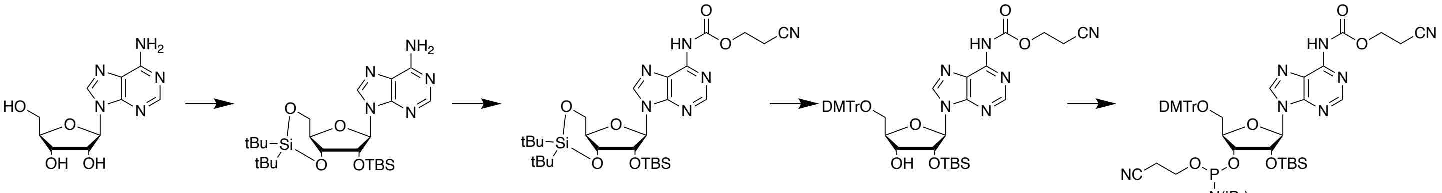


**Adenosine** **1** **2** **3** **4**

**Scheme S1.** Synthesis of *N*-ceoc protected adenosine phosphoramidite.

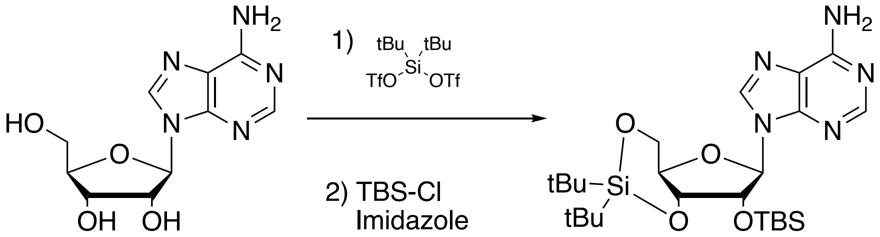


Adenosine (3.00 g, 11.2 mmol) was dissolved in DMF (56 mL) and cooled to 0 °C. Di-tert-butylsilyl bis(trifluoromethanesulfonate) (4.40 mL, 13.5 mmol) was added dropwise to the stirred DMF solution. After 40 min at 0 °C, imidazole (3.81 g, 56.0 mmol) was added and the reaction mixture was allowed to warm to 23 °C. TBS-Cl (2.03 g, 13.5 mmol) was added in one portion and the stirred reaction mixture was heated to 60 °C for 18 h. DMF was removed under reduced pressure. Water (100 mL) was added to the resultant residue to precipitate a light pink solid. The solid was collect by vacuum filtration. The solid was recrystallized in dichloromethane to yield 1.81 g of crystalline **1**. The mother liquor was condensed under reduced pressure and the resultant solid was purified by silica flash chromatography (1:1 to 3:1 ethyl acetate: hexanes) to yield an additional 2.48 g of **1** as a white solid (total yield: 4.29 g, 73% yield). $^1$H NMR (500 MHz, DMSO) δ 8.34 (s, 1H), 8.12 (s, 1H), 7.34 (s, 2H), 5.95 (s, 1H), 4.76 – 4.70 (m, 1H), 4.69 – 4.64 (m, 1H), 4.38 – 4.35 (m, 1H), 4.00 (d, *J* = 6.1 Hz, 2H), 1.08 (s, 9H), 1.01 (s, 9H), 0.86 (s, 9H), 0.09 (s, 3H), 0.07 (s, 3H). $^{13}$C NMR (126 MHz, DMSO) δ 156.06, 152.64, 148.77, 139.79, 119.10, 91.02, 75.21, 74.81, 74.08, 67.02, 27.56, 27.32, 26.82, 25.80, 25.69, 22.21, 19.96, 18.05, -3.20, -4.59, -5.18. HRMS (ESI/Q-TOF) calcd for $C_{24}H_{44}N_5O_4Si_2^+$ [M + H]$^+$ 522.2926, found 522.2934

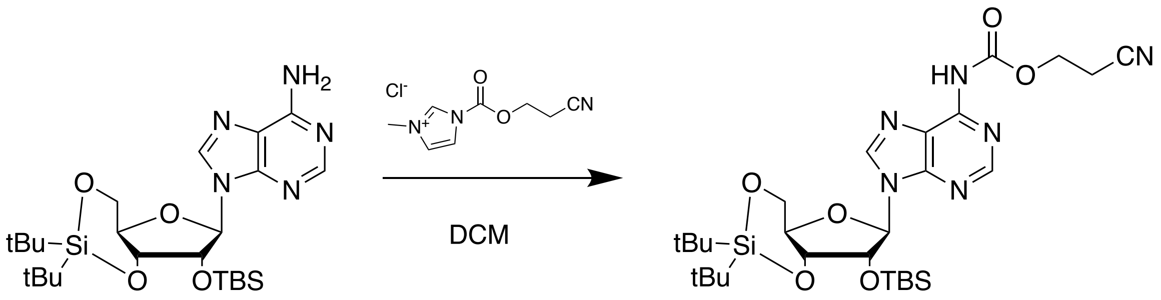


Protected adenosine **1** (4.00 g, 7.67 mmol) was dissolved in dichloromethane (20 mL). Ceoc-carbonyl-*N*-methylimidazolium chloride (2.06 g, 9.59 mmol) was added in one portion. After stirring for 18 h, more ceoc-carbonyl-*N*-methylimidazolium chloride was added (0.825 g, 3.84 mmol) and stirred an additional 24 h. The reaction mixture was diluted with DCM (200 mL) transferred to a separatory funnel, and washed with water (200 mL), sat'd aq. $NaHCO_3$ (200 mL), and brine (200 mL). The organic layer was dried over anhydrous $Na_2SO_4$, filtered, and condensed under reduced pressure. The crude product was purified by silica flash chromatography (6:4 ethyl acetate:hexanes) to yield *N*-ceoc protected **2** as a white solid (4.53 g, 95% yield). Characterization: $^1$H NMR (500 MHz, DMSO) δ 10.83 (s, 1H), 8.65 (s, 1H), 8.63 (s, 1H), 6.06 (s, 1H), 4.71 (d, *J* = 5.0 Hz, 1H), 4.68 (td, *J* = 8.1, 7.6, 4.9 Hz, 1H), 4.41 – 4.35 (m, 1H), 4.32 (t, *J* = 6.0 Hz, 2H), 4.30 (t, *J* = 5.9 Hz, 1H), 4.09 – 4.03 (m, 2H), 3.32 (s, 2H), 2.95 (td, *J* = 5.9, 4.1 Hz, 3H), 1.07 (s, 9H), 1.01 (s, 10H), 0.88 (s, 9H), 0.11 (s, 3H), 0.08 (s, 3H). $^{13}$C NMR (126 MHz, DMSO) δ 153.51, 151.70, 151.17, 149.64, 142.97, 124.05, 118.52, 118.29, 91.16, 75.18, 74.78, 74.22, 66.95, 62.76, 59.95, 27.31, 26.81, 25.70, 22.21, 19.96, 18.00, 17.66, 17.40, -4.58, -5.17. HRMS (ESI/Q-TOF) calcd for $C_{28}H_{47}N_6O_6Si_2^+$ [M + H]$^+$ 619.3090, found 619.3189.

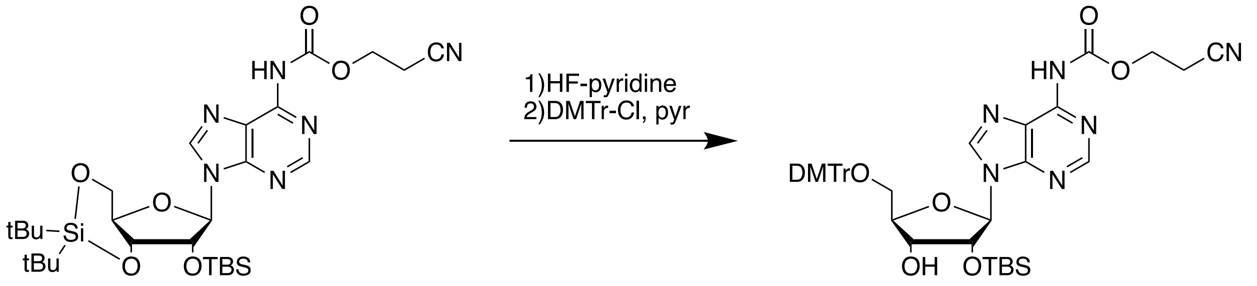


HF-pyridine (0.67 mL, 26 mmol) was added to pyridine (4.20 mL, 52.0 mmol) in a 15 mL plastic conical tube at 0 °C. Compound **2** (4.00 g, 6.46 mmol) was suspended in DCM (32 mL) in a 50 mL plastic conical tube and cooled to 0 °C. The HF-pyridine solution was added dropwise to the DCM suspension of **2** at 0 °C with stirring. After 90 minutes, the reaction was complete as evidenced by TLC (4:6 hexanes:ethyl acetate) analysis. The reaction mixture was diluted with DCM (150 mL), and washed with 150 mL each of water, saturated aqueous sodium bicarbonate, and brine. The organic layer was collected and dried over anhydrous $Na_2SO_4$, filtered, and then condensed under reduced pressure. The crude diol was dissolved in pyridine (13 mL) and cooled to 0 °C. Dimethoxytrityl chloride (2.41 g, 7.11 mmol) was added to the pyridine solution at 0 °C with stirring. After 10 minutes, the seal reaction vessel was transferred to a 4 °C refrigerator for 16 h. To quench the reaction, methanol (3 mL) was added and the reaction mixture was stirred at 23 °C for 5 minutes after which, all volatiles were removed under reduced pressure. The crude product was purified by silica flash chromatography (3:7 to 1:1 ethyl acetate: hexanes) to yield **3** (3.82 g, 76% yield). Characterization: $^1$H NMR (500 MHz, DMSO) δ 10.82 (s, 1H), 8.59 (s, 1H), 8.57 (s, 1H), 7.39 (dt, *J* = 6.4, 1.3 Hz, 2H), 7.30 – 7.17 (m, 8H), 6.88 – 6.81 (m, 4H), 6.06 (d, *J* = 4.8 Hz, 1H), 5.20 (d, *J* = 5.9 Hz, 1H), 4.87 (t, *J* = 4.9 Hz, 1H), 4.32 (t, *J* = 6.0 Hz, 2H), 4.28 (q, *J* = 5.2 Hz, 1H), 4.13 (q, *J* = 4.5 Hz, 1H), 3.73 (s, 6H), 3.32 – 3.25 (m, 2H), 2.94 (t, *J* = 6.0 Hz, 2H), 0.75 (s, 9H), -0.04 (s, 3H), -0.14 (s, 3H). $^{13}$C NMR (126 MHz, DMSO) δ 158.07, 151.65, 151.59, 149.61, 144.85, 142.94, 135.52,

135.40, 129.72, 127.79, 127.65, 126.69, 124.01, 118.53, 113.15, 88.23, 85.55, 83.52, 74.84, 70.14, 63.38, 59.93, 55.02, 25.55, 17.82, 17.66, -4.83, -5.29. HRMS (ESI/Q-TOF) calcd for $C_{41}H_{49}N_6O_8Si^+$ $[M + H]^+$ 781.3376, found 781.3433.

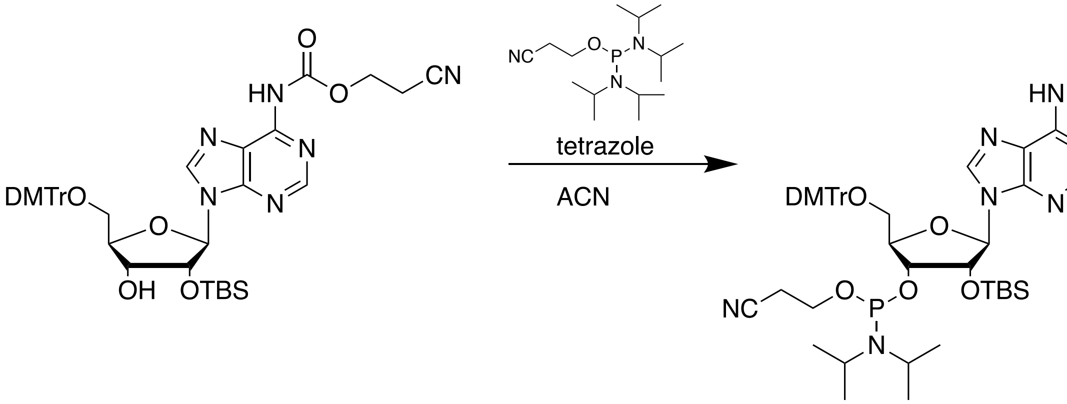


Compound **3** (3.82 g, 4.89 mmol) was dissolved in acetonitrile (30 mL) and a solution of tetrazole in acetonitrile (0.45 M, 10.9 mL, 4.89 mmol) was added. 2-cyanoethyl bis(N,N-diisopropyl) phosphordiamidite (2.33 mL, 7.34 mmol) was added dropwise and the reaction mixture was stirred at 23 °C for 12 h. Monitoring by reversed-phase HPLC indicated complete conversion. Acetonitrile was removed under reduced pressure and the resultant residue was dissolved in ethyl acetate (150 mL) and transferred to a separatory funnel. The organic layer was washed with sat'd aq. $NaHCO_3$ (150 mL), then dried over anhydrous $Na_2SO_4$, filtered, and then condensed under reduced pressure. Purified by $C_{18}$ flash chromatography (solvent A: 20 mM triethylammonium acetate in water,pH 7; solvent B: ACN; 70% B for 1 column volumes (CV), 70 to 100% B over 5 CVs, 100% B for 8 CVs). Fractions containing amidite **4** were pooled and condensed under reduced pressure. To remove water introduced during purification, the resultant solid was dissolved in ethyl acetate (50 mL) and dried over anhydrous $Na_2SO_4$, filtered, and then condensed under reduced pressure to yield **4** as a white foam (2.31 g, 48% yield, a mixture of diastereomers). $^1$H NMR (500 MHz, $CDCl_3$) δ 8.66 (d, *J* = 10.4 Hz, 1H), 8.49 (s, 1H), 8.22 (dd, *J* = 14.7, 1.3 Hz, 1H), 7.46 (tt, *J* = 8.0, 1.4 Hz, 2H), 7.40 – 7.31 (m, 4H), 7.28 (dd, *J* = 7.0, 4.4 Hz, 1H), 7.28 – 7.17 (m, 1H), 6.86 – 6.78 (m, 4H), 6.08 (d, *J* = 6.3 Hz, 0H), 6.03 (d, *J* = 6.0 Hz, 0H), 5.05 (ddd, *J* = 10.4, 6.1, 4.5 Hz, 1H), 4.46 (tdd, *J* = 8.7, 6.0, 3.1 Hz, 2H), 4.45 – 4.33 (m, 1H), 4.12 (q, *J* = 7.1 Hz, 1H), 3.78 (dd, *J* = 3.1, 1.4 Hz, 6H), 3.72 – 3.53 (m, 3H), 3.34 (td, *J* = 11.1, 3.8 Hz, 1H), 2.81 (td, *J* = 6.4, 2.1 Hz, 2H), 2.69 – 2.60 (m, 1H), 2.31 (td, *J* = 6.5, 4.3 Hz, 1H), 2.04 (s, 1H), 1.72 (s, 3H), 1.26 (t, *J* = 7.1 Hz, 1H), 1.22 – 1.14 (m, 8H), 1.05 (d, *J* = 6.7 Hz, 3H), 0.75 (s, 8H), -0.03 (s, 1H), -0.05 (s, 1H), -0.21 (s, 2H), -0.23 (s, 1H). $^{13}$C NMR (126 MHz, $CDCl_3$) δ 158.71, 152.83, 151.57, 150.29, 148.98, 144.69, 144.57, 142.38, 142.36, 135.83, 135.79, 135.65, 135.61, 130.30, 130.25, 130.21, 130.20, 128.36, 128.25, 128.04, 128.02, 127.11, 122.71, 122.67, 117.73, 117.41, 116.76, 113.33, 113.31, 113.29, 88.66, 88.44, 86.91, 86.77, 84.37, 84.08, 84.05, 75.26, 74.71, 74.67, 73.49, 73.41, 72.86, 72.74, 63.36, 63.20, 60.51, 60.23, 58.99, 58.85, 57.81, 57.65, 55.38, 55.35, 47.52, 43.59, 43.49, 43.12, 43.02, 25.73, 25.69, 24.90, 24.81, 24.75, 24.68, 20.61, 20.56, 20.23, 20.18, 19.56, 18.30, 18.03, 17.97, 14.32, -4.56, -4.58, -5.04. HRMS (ESI/Q-TOF) calcd for $C_{50}H_{66}N_8O_9PSi^+$ $[M + H]^+$ 981.4454, found 981.9879.

*Cytidine*

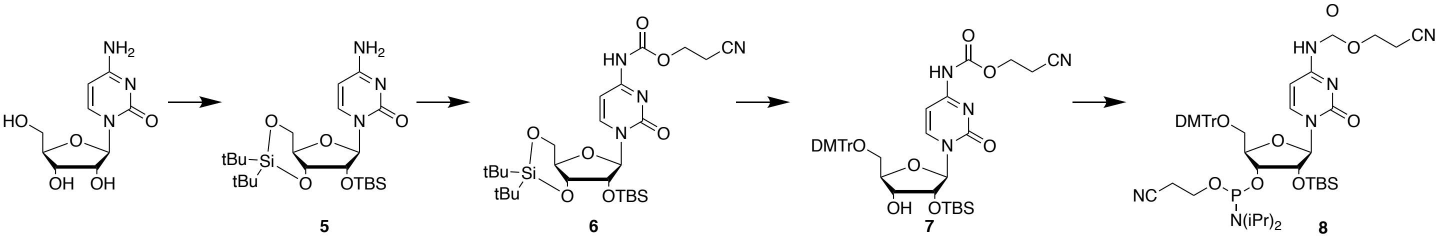


**Scheme S2**. Synthesis of *N*-ceoc protected cytidine phosphoramidite.

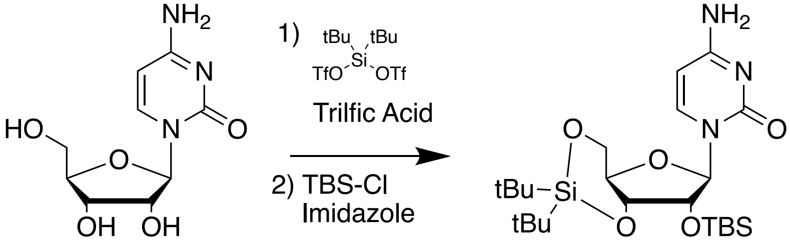


Cytosine (4.00 g, 16.4 mmol) was suspended in DMF (41 mL) and cooled to 0 °C. Triflic acid (1.45 mL, 16.4 mmol) was added dropwise resulting in a homogenous solution. Di-tert-butylsilyl bis(trifluoromethanesulfonate) (6.42 mL, 19.7 mmol) was added dropwise to the stirred DMF solution. After 45 min at 0 °C, imidazole (6.7 g, 98.4 mmol) was added and the reaction mixture was allowed to warm to 23 °C. TBS-Cl (2.97 g, 19.7 mmol) was added in one portion and the stirred reaction mixture was heated to 60 °C for 2h. DMF was removed under reduced pressure. The crude reaction mixture was dissolved in ethyl acetate (200 mL), transferred to a separatory funnel, and washed with water (2 x 200 mL) and brine (200 mL). The organic layer was dried over anhydrous $Na_2SO_4$, filtered, and then condensed under reduced pressure. Purification by silica flash chromatography yielded ribose-protected cytosine **5** as a white solid (6.25 g, 77% yield). $^1$H NMR (500 MHz, DMSO) δ 7.54 (d, *J* = 7.5 Hz, 1H), 7.28 (d, *J* = 18.1 Hz, 2H), 5.74 (d, *J* = 7.4 Hz, 1H), 5.66 (s, 1H), 4.36 (dd, *J* = 9.0, 5.0 Hz, 1H), 4.27 (d, *J* = 4.8 Hz, 1H), 4.07 – 3.95 (m, 2H), 3.90 (td, *J* = 10.0, 4.9 Hz, 1H), 3.17 (s, 1H), 1.02 (s, 9H), 0.98 (s, 9H), 0.87 (s, 9H), 0.12 (s, 3H), 0.08 (s, 3H). $^{13}$C NMR (126 MHz, DMSO) δ 165.64, 154.54, 141.90, 94.48, 93.88, 75.31, 74.61, 73.70, 66.94, 48.59, 27.32, 26.79, 25.72, 22.17, 19.93, 18.04, -4.60, -5.10. HRMS (ESI/Q-TOF) calcd for $C_{23}H_{43}N_3O_5Si_2Na^+$ $[M + Na]^+$ 520.2633, found 520.2632.

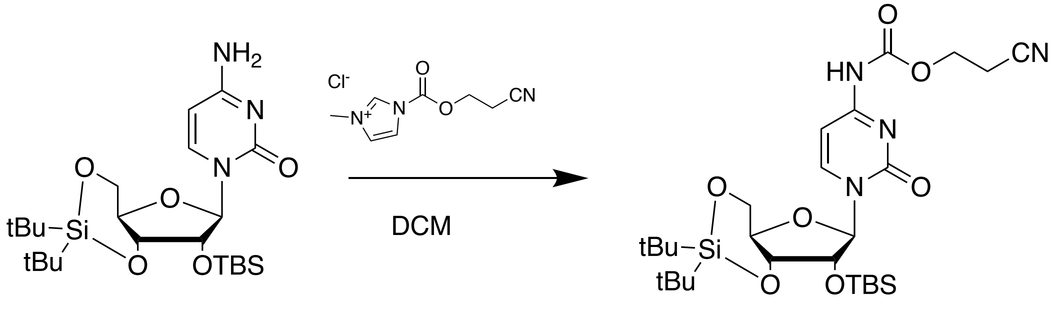


Protected cytosine **5** (4.00 g, 8.04 mmol) was dissolved in dichloromethane (20 mL). Ceoc-carbonyl-*N*-methylimidazolium chloride (2.16 g, 10.0 mmol) was added in one portion. After stirring for 18 h, more ceoc-carbonyl-*N*-methylimidazolium chloride was added (0.864 g, 4.02 mmol) and stirred an additional 24 h. The reaction mixture was diluted with DCM (200 mL) transferred to a separatory funnel, and washed with water (200 mL), sat'd aq. $NaHCO_3$ (200 mL), and brine (200 mL). The organic layer was dried over anhydrous

$Na_2SO_4$, filtered, and condensed under reduced pressure. The crude product was purified by silica flash chromatography (6:4 ethyl acetate:hexanes) to yield *N*-ceoc protected **6** as a white solid (4.78 g, quantitative yield). $^1$H NMR (500 MHz, DMSO) δ 10.94 (s, 1H), 8.00 (d, *J* = 7.6 Hz, 1H), 7.04 (d, *J* = 7.6 Hz, 1H), 5.69 (s, 1H), 4.41 (dd, *J* = 9.1, 4.8 Hz, 1H), 4.35 – 4.23 (m, 4H), 4.14 (t, *J* = 9.6 Hz, 1H), 4.02 (td, *J* = 9.8, 4.8 Hz, 1H), 3.98 (dd, *J* = 9.6, 4.4 Hz, 1H), 2.98 – 2.89 (m, 3H), 1.02 (s, 9H), 0.98 (s, 11H), 0.90 (s, 9H), 0.17 (s, 3H), 0.11 (s, 3H). $^{13}$C NMR (126 MHz, DMSO) δ 162.88, 154.62, 153.84, 153.51, 145.06, 118.40, 94.85, 93.54, 74.89, 74.69, 74.20, 66.79, 62.75, 62.39, 60.12, 54.93, 27.33, 26.76, 25.72, 22.15, 19.93, 17.96, 17.59, -4.56, -5.01. HRMS (ESI/Q-TOF) calcd for $C_{27}H_{47}N_4O_7Si_2^+$ [M + H]$^+$ 595.2978, found 595.2977.

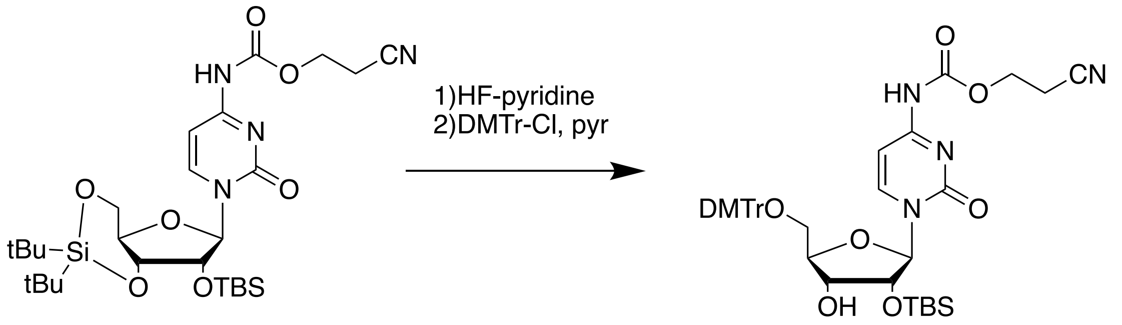


HF-pyridine (0.79 mL, 30.3 mmol) was added to pyridine (4.91 mL, 60.9 mmol) in a 15 mL plastic conical tube at 0 °C. Compound **6** (4.50 g, 7.56 mmol) was suspended in DCM (38 mL) in a 50 mL plastic conical tube and cooled to 0 °C. The HF-pyridine solution was added dropwise to the DCM suspension of **6** at 0 °C with stirring. After 90 minutes, the reaction was complete as evidenced by TLC (4:6 hexanes:ethyl acetate) analysis. The reaction mixture was diluted with DCM (150 mL), and washed with 150 mL each of water, saturated aqueous sodium bicarbonate, and brine. The organic layer was collected and dried over anhydrous $Na_2SO_4$, filtered, and then condensed under reduced pressure. The crude diol was dissolved in pyridine (15 mL) and cooled to 0 °C. Dimethoxytrityl chloride (2.818 g, 8.32 mmol) was added to the pyridine solution at 0 °C with stirring. After 10 minutes, the seal reaction vessel was transferred to a 4 °C refrigerator for 16 h. To quench the reaction, methanol (3 mL) was added and the reaction mixture was stirred at 23 °C for 5 minutes after which, all volatiles were removed under reduced pressure. The crude product was purified by silica flash chromatography (1:0 to 95:5 dichloromethane: methanol) to yield **7** (3.59 g, 63% yield). $^1$H NMR (500 MHz, DMSO) δ 10.92 (s, 1H), 8.31 (d, *J* = 7.5 Hz, 1H), 7.40 (d, *J* = 7.6 Hz, 2H), 7.33 (t, *J* = 7.7 Hz, 2H), 7.26 (dd, *J* = 8.7, 1.8 Hz, 5H), 6.91 (dd, *J* = 9.0, 1.5 Hz, 4H), 6.82 – 6.75 (m, 1H), 5.72 (d, *J* = 1.8 Hz, 1H), 5.13 (d, *J* = 6.2 Hz, 1H), 4.30 (t, *J* = 6.0 Hz, 2H), 4.17 (ddd, *J* = 7.9, 6.1, 4.4 Hz, 1H), 4.13 (dd, *J* = 4.5, 1.8 Hz, 1H), 4.06 (dt, *J* = 8.1, 3.1 Hz, 1H), 3.75 (s, 6H), 3.35 (qd, *J* = 11.3, 3.6 Hz, 2H), 2.93 (t, *J* = 6.0 Hz, 2H), 0.88 (s, 9H), 0.12 (s, 3H), 0.08 (s, 3H). $^{13}$C NMR (126 MHz, DMSO) δ 162.71, 158.17, 152.69, 144.37, 144.18, 135.46, 135.15, 129.79, 129.68, 127.94, 127.80, 126.88, 118.42, 113.28, 94.24, 90.73, 86.00, 81.59, 76.33, 68.21, 61.73, 60.11, 55.02, 25.74, 17.98, 17.57, -4.75, -4.89. HRMS (ESI/Q-TOF) calcd for $C_{40}H_{49}N_4O_9Si^+$ [M + H]$^+$ 757.3263, found 757.3283.

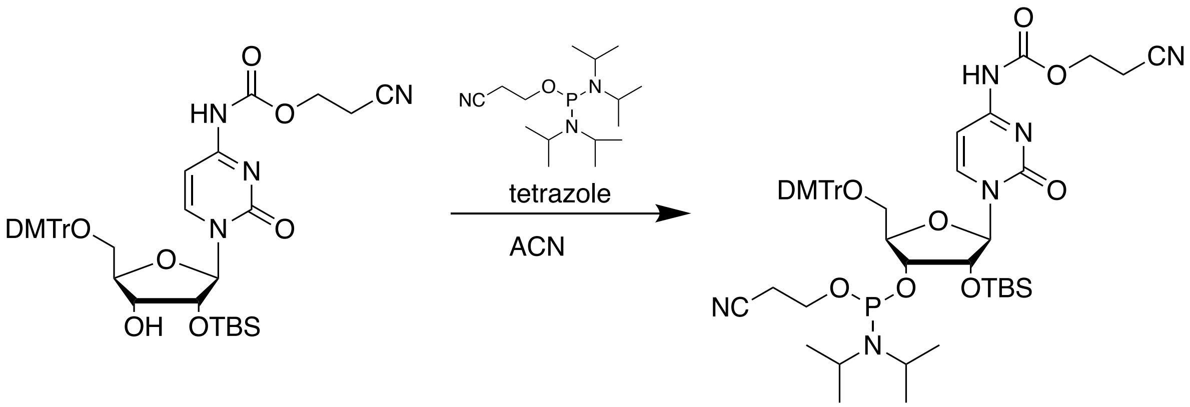

Compound **7** (3.57 g, 4.71 mmol) was dissolved in acetonitrile (28 mL) and a solution of tetrazole in acetonitrile (0.45 M, 10.5 mL, 4.71 mmol) was added. 2-cyanoethyl bis(N,N-diisopropyl) phosphordiamidite (2.24 mL, 7.07 mmol) was added dropwise and the reaction mixture was stirred at 23 °C for 12 h. Monitoring by reversed-phase HPLC indicated complete conversion. Acetonitrile was removed under reduced pressure and the resultant residue was dissolved in ethyl acetate (150 mL) and transferred to a separatory funnel. The organic layer was washed with sat'd aq. $NaHCO_3$ (150 mL), then dried over anhydrous $Na_2SO_4$, filtered, and then condensed under reduced pressure. Purified by $C_{18}$ flash chromatography (solvent A: 20 mM triethylammonium acetate in water,pH 7; solvent B: ACN; 70% B for 1 column volumes (CV), 70 to 100% B over 5 CVs, 100% B for 8 CVs). Fractions containing amidite **8** were pooled and condensed under reduced pressure. To remove water introduced during purification, the resultant solid was dissolved in ethyl acetate (50 mL) and dried over anhydrous $Na_2SO_4$, filtered, and then condensed under reduced pressure to yield **8** as a white foam (2.04 g, 45% yield, a mixture of diastereomers). $^1$H NMR (400 MHz, $CDCl_3$) δ 7.93 (d, *J* = 7.4 Hz, 1H), 7.79 (d, *J* = 7.4 Hz, 1H), 7.23 – 7.16 (m, 2H), 7.16 – 7.12 (m, 2H), 7.08 (d, *J* = 8.8 Hz, 4H), 7.06 – 6.96 (m, 10H), 6.57 (ddd, *J* = 8.9, 5.7, 2.3 Hz, 9H), 5.73 (d, *J* = 3.3 Hz, 1H), 5.60 (d, *J* = 1.8 Hz, 1H), 5.22 (s, 2H), 4.97 (dd, *J* = 15.4, 7.4 Hz, 2H), 4.10 (t, *J* = 3.5 Hz, 1H), 4.07 – 3.93 (m, 6H), 3.74 – 3.58 (m, 1H), 3.56 – 3.48 (m, 13H), 3.48 – 3.20 (m, 7H), 3.12 (ddd, *J* = 16.2, 10.9, 2.5 Hz, 2H), 2.54 (d, *J* = 4.9 Hz, 1H), 2.35 (td, *J* = 6.4, 1.4 Hz, 2H), 2.27 (q, *J* = 7.1 Hz, 8H), 2.12 (t, *J* = 6.6 Hz, 2H), 1.50 (s, 5H), 0.91 – 0.82 (m, 21H), 0.77 (t, *J* = 7.2 Hz, 12H), 0.70 (d, *J* = 6.8 Hz, 6H), 0.64 (d, *J* = 4.5 Hz, 19H), -0.05 (d, *J* = 8.5 Hz, 6H), -0.14 (s, 6H). $^{13}$C NMR (126 MHz, $CDCl_3$) δ 158.83, 145.12, 144.22, 135.55, 135.38, 130.42, 130.38, 130.36, 128.55, 128.27, 128.12, 128.07, 127.33, 117.61, 117.46, 116.58, 113.52, 113.38, 113.34, 113.32, 94.88, 87.34, 87.24, 81.51, 75.96, 75.39, 75.37, 61.05, 60.51, 60.11, 58.36, 58.20, 55.40, 55.36, 55.33, 47.11, 43.36, 43.26, 43.19, 43.09, 25.97, 25.91, 25.06 – 24.55 (m), 21.17, 20.55, 20.50, 20.33, 20.27, 19.16, 19.09, 18.16, 18.09, 14.32, -4.25, -4.97.

<u>*Dihydrouridine*</u>

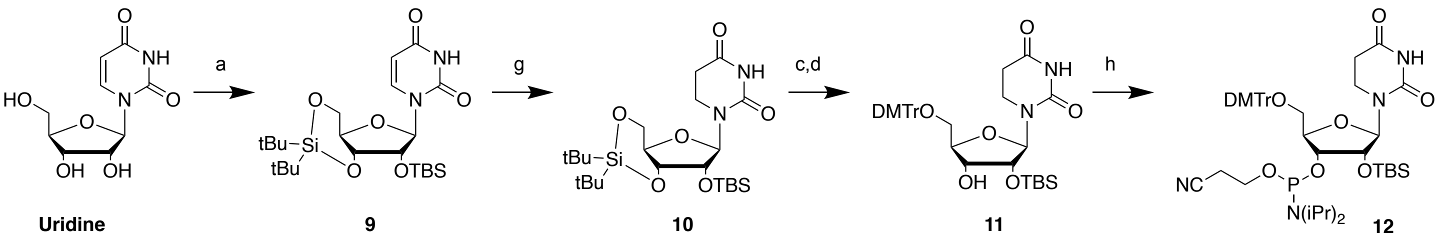


**Scheme S3**. Synthesis of dihydrouridine phosphoramidite.

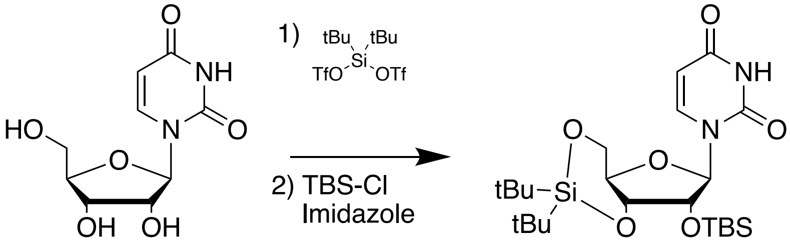


Uridine (3.00 g, 12.3 mmol) was dissolved in anhydrous DMF (15 mL) and cooled to 0 °C. Di-tert-butylsilyl bis(trifluoromethanesulfonate) (4.81 mL, 14.7 mmol) was added dropwise to the stirred DMF solution. After 45 min at 0 °C, imidazole (4.19 g, 61.5 mmol) was added and the reaction mixture was allowed to warm to 23 °C. TBS-Cl (2.22 g, 14.7 mmol) was added in one portion and the reaction mixture was stirred at 23 °C for 16h. DMF was removed under reduced pressure. The crude reaction mixture was dissolved in ethyl acetate (200 mL), transferred to a separatory funnel, and washed with water (2 x 200 mL) and brine (200 mL). The organic layer was dried over anhydrous $Na_2SO_4$, filtered, and then condensed under reduced pressure. Purification by silica flash chromatography (3:7 ethyl acetate:hexanes) yielded ribose-protected uridine **9** as a white solid (4.55 g, 74% yield). $^1$H NMR (500 MHz, DMSO) δ 11.39 (d, *J* = 2.3 Hz, 1H), 7.59 (d, *J* = 8.1 Hz, 1H), 5.65 (s, 1H), 5.61 (dd, *J* = 8.1, 2.2 Hz, 1H), 4.37 (dd, *J* = 4.7, 3.4 Hz, 2H), 4.03 (ddd, *J* = 15.8, 10.1, 7.1 Hz, 2H), 3.90 (td, *J* = 10.0, 5.0 Hz, 1H), 1.03 (s, 9H), 0.98 (s, 10H), 0.88 (s, 10H), 0.12 (s, 3H), 0.08 (s, 3H). $^{13}$C NMR (126 MHz, DMSO) δ 163.07, 150.06, 141.13, 101.98, 92.94, 75.15, 74.36, 73.87, 66.78, 27.30, 26.77, 25.69, 22.15, 19.93, 18.02, -4.55, -5.23. HRMS (ESI/Q-TOF) calcd for $C_{23}H_{43}N_2O_6Si_2^+$ [M + H]$^+$ 499.2654, found 499.2687.

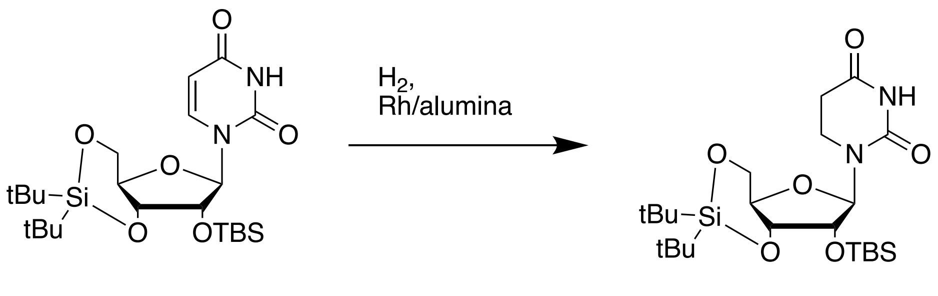


Ribose-protected uridine **9** (4.00 g, 8.02 mmol) was dissolved in MeOH (120 mL) and Rh/alumina (5% wt/wt loading, 0.80 g) was added. The solution was sparged with $H_2$ for 5 min then the solution was stirred vigorously under $H_2$ for 16 h at 23 °C. The catalyst was removed by first vacuum filtration through paper filter, and then by filtrations through a 0.45 mm PTFE filter. MeOH was removed under reduced pressure and the crude product was purified by silica flash chromatography (20 to 30% ethyl acetate in hexanes) to yield ribose-protected dihydrouridine **10** as a white solid (2.512 g, 63% yield). $^1$H NMR (500 MHz, DMSO) δ 10.33 (s, 1H), 5.66 (d, *J* = 1.2 Hz, 1H), 4.33 (dd, *J* = 9.2, 4.9 Hz, 1H), 4.31 (dd, *J* = 5.4, 1.2 Hz, 1H), 3.89 – 3.82 (m, 2H), 3.75 (td, *J* = 10.2, 4.9 Hz, 1H), 3.40 – 3.25 (m, 2H), 2.55 – 2.46 (m, 1H), 1.03 (s, 9H), 0.98 (s, 9H), 0.88 (s, 9H), 0.09 (d, *J* = 17.3 Hz, 6H). $^{13}$C NMR (126 MHz, DMSO) δ 170.23, 152.41, 92.75, 76.29, 73.34, 72.81, 67.13, 37.12, 30.88, 27.27, 26.81, 25.71, 22.24, 19.94, 18.09, -4.57, -5.25. HRMS (ESI/Q-TOF) calcd for $C_{23}H_{45}N_2O_6Si_2^+$ [M + H]$^+$ 501.2811, found 501.2826.

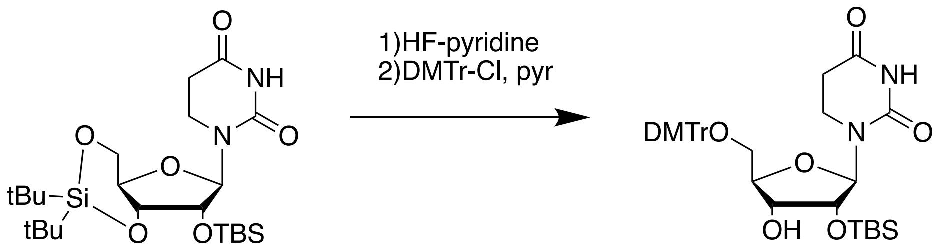


HF-pyridine (0.48 mL, 18.4 mmol) was added to pyridine (2.98 mL, 36.9 mmol) in a 15 mL plastic conical tube at 0 °C. Compound **10** (2.30 g, 4.59 mmol) was suspended in DCM (23 mL) in a 50 mL plastic conical tube and cooled to 0 °C. The HF-pyridine solution was added dropwise to the DCM suspension of **10** at 0 °C with stirring. After 90 minutes, the reaction was complete as evidenced by TLC (7:3 hexanes:ethyl acetate) analysis. The reaction mixture was diluted with DCM (150 mL), and washed with 150 mL each of water, saturated aqueous sodium bicarbonate, and brine. The organic layer was collected and dried over anhydrous $Na_2SO_4$, filtered, and then condensed under reduced pressure. The crude diol was dissolved in pyridine (9 mL) and cooled to 0 °C. Dimethoxytrityl chloride (1.711 g, 5.05 mmol) was added to the pyridine solution at 0 °C with stirring. After 10 minutes, the seal reaction vessel was transferred to a 4 °C refrigerator for 16 h. To quench the reaction, methanol (3 mL) was added and the reaction mixture was stirred at 23 °C for 5 minutes after which, all volatiles were removed under reduced pressure. The crude product was purified by silica flash chromatography (30 to 50% ethyl acetate in hexanes) to yield **11** (2.319 g, 76% yield). $^1$H NMR (500 MHz, DMSO) δ 10.31 (s, 1H), 7.38 (d, *J* = 8.4 Hz, 2H), 7.32 (t, *J* = 7.7 Hz, 2H), 7.28 – 7.19 (m, 5H), 6.90 (d, *J* = 8.5

Hz, 4H), 5.74 (d, *J* = 4.7 Hz, 1H), 4.82 (d, *J* = 6.4 Hz, 1H), 4.12 (t, *J* = 5.0 Hz, 1H), 3.92 – 3.82 (m, 2H), 3.74 (s, 6H), 3.44 (ddd, *J* = 12.8, 7.2, 5.6 Hz, 1H), 3.35 – 3.27 (m, 1H), 3.21 – 3.10 (m, 2H), 2.50 – 2.45 (m, 1H), 2.40 (ddd, *J* = 16.6, 7.3, 5.3 Hz, 1H), 0.87 (s, 9H), 0.09 (s, 6H). $^{13}$C NMR (126 MHz, DMSO) δ 170.16, 158.09, 152.74, 149.61, 144.86, 136.12, 135.47, 135.37, 129.72, 127.85, 127.64, 126.71, 123.90, 113.21, 88.19, 85.52, 81.65, 73.12, 70.13, 63.57, 55.04, 35.91, 30.79, 25.70, 17.99, -4.61, -5.04. HRMS (ESI/Q-TOF) calcd for $C_{36}H_{46}N_2O_8SiNa^+$ [M + Na]$^+$ 685.2916, found 685.2919.

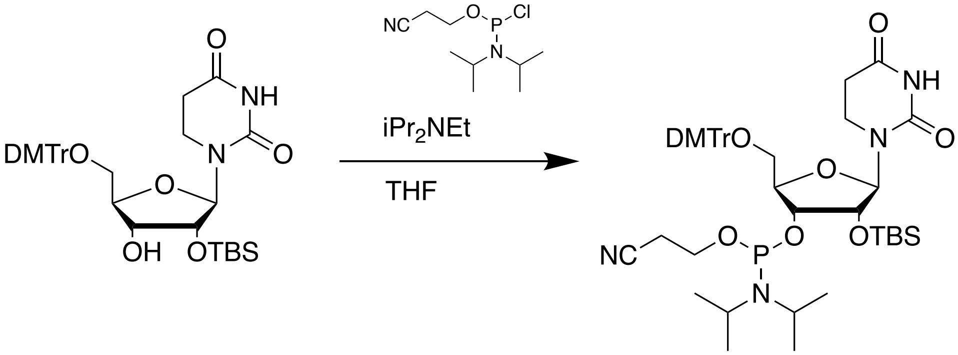


To a solution of dihydrouridine **11** (2.20 g, 3.32 mmol) in THF (30 mL) was added $iPr_2NEt$ (2.32 mL, 13.3 mmol) immediately followed by addition of 2-cyanoethyl *N,N*-diisopropyl phosphoamidochloridite (1.48 mL, 6.64 mmol). After 20 h, monitoring by reversed-phase HPLC indicated complete conversion. The reaction mixture was diluted with ethyl acetate (150 mL) and transferred to a separatory funnel. The organic layer was washed with water (200 mL), sat'd aq. $NaHCO_3$ (200 mL), then dried over anhydrous $Na_2SO_4$, filtered, and then condensed under reduced pressure. Purified by $C_{18}$ flash chromatography (solvent A: 20 mM triethylammonium acetate in water,pH 7; solvent B: ACN; 70% B for 1 column volumes (CV), 70 to 100% B over 5 CVs, 100% B for 8 CVs). Fractions containing amidite **12** were pooled and condensed under reduced pressure. To remove water introduced during purification, the resultant solid was dissolved in ethyl acetate (50 mL) and dried over anhydrous $Na_2SO_4$, filtered, and then condensed under reduced pressure to yield **12** as a white foam (2.263 g, 79% yield, a mixture of diastereomers). $^1$H NMR (500 MHz, DMSO) δ 10.35 (d, *J* = 8.0 Hz, 1H), 7.43 – 7.35 (m, 2H), 7.32 (td, *J* = 7.6, 5.5 Hz, 2H), 7.24 (ddt, *J* = 11.5, 9.3, 3.1 Hz, 5H), 6.90 (t, *J* = 8.4 Hz, 4H), 5.75 (dd, *J* = 10.5, 5.8 Hz, 1H), 4.28 (dt, *J* = 13.7, 5.4 Hz, 1H), 4.13 – 4.06 (m, 1H), 3.99 (q, *J* = 3.7 Hz, 1H), 3.73 (d, *J* = 1.9 Hz, 7H), 3.62 – 3.42 (m, 4H), 3.22 (dd, *J* = 10.7, 2.8 Hz, 1H), 3.13 (ddd, *J* = 10.8, 7.4, 4.3 Hz, 1H), 2.76 (dt, *J* = 6.2, 4.9 Hz, 1H), 2.57 – 2.45 (m, 1H), 2.40 (ddt, *J* = 17.0, 7.4, 5.3 Hz, 1H), 1.13 – 1.07 (m, 9H), 0.94 (d, *J* = 6.8 Hz, 3H), 0.87 (d, *J* = 9.5 Hz, 9H), 0.11 (d, *J* = 6.6 Hz, 3H), 0.08 (d, *J* = 1.7 Hz, 3H). $^{13}$C NMR (126 MHz, DMSO) δ 170.05, 170.01, 158.16, 158.13, 152.93, 152.86, 144.76, 144.63, 135.31, 135.27, 135.18, 135.03, 129.73, 129.71, 129.66, 127.86, 127.63, 127.54, 126.78, 126.75, 118.85, 118.64, 113.22, 113.20, 87.53, 87.51, 85.80, 85.75, 81.26, 80.95, 71.93, 71.53, 63.21, 59.74, 58.56, 58.42, 57.61, 57.44, 55.05, 42.77, 42.67, 42.33, 42.23, 35.99, 35.87, 30.75, 30.69, 25.63, 25.58, 24.45, 24.40, 24.33, 24.27, 24.22, 24.17, 19.90, 19.85, 19.74, 19.69, 17.75, 17.70, -4.45, -4.48, -4.57, -4.59, -5.02, -5.04.

*Guanosine*

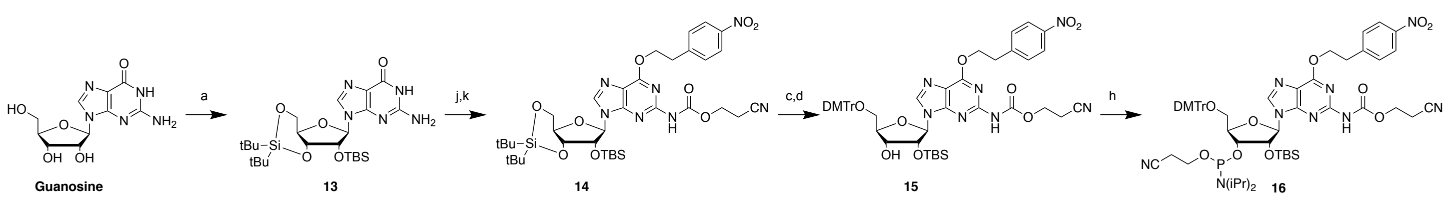

**Scheme S4**. Synthesis of *N*-ceoc protected guanosine phosphoramidite.

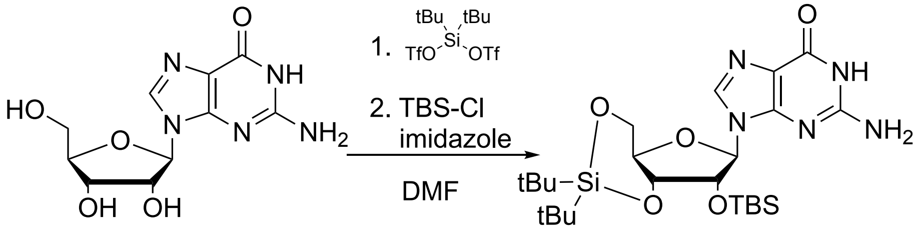

Guanosine (12.75 g, 45.0 mmol) was suspended in DMF (90 mL) and cooled to 0 °C. Di-tert-butylsilyl bis(trifluoromethanesulfonate) (16.15 mL, 49.5 mmol) was added dropwise to the stirred DMF solution. After 45 min at 0 °C, imidazole (15.31 g, 225.0 mmol) was added and the reaction mixture was allowed to warm to 23 °C. TBS-Cl (8.14 g, 54.0 mmol) was added in one portion and the stirred reaction mixture was heated to 60 °C for 2h. DMF was removed under reduced pressure and the resulting residue was recrystallized from MeOH to yield ribose-protected guanosine **13** as a white solid (12.10 g, 50% yield). $^1$H NMR (500 MHz, DMSO) δ 10.66 (s, 1H), 7.91 (s, 1H), 6.36 (s, 2H), 5.72 (d, *J* = 1.1 Hz, 1H), 4.59 – 4.55 (m, 1H), 4.34 (dd, *J* = 8.3, 4.2 Hz, 1H), 4.28 (dd, *J* = 9.0, 5.2 Hz, 1H), 4.01 – 3.90 (m, 2H), 1.06 (s, 10H), 1.00 (s, 10H), 0.86 (s, 9H), 0.09 (s, 3H), 0.07 (s, 3H). $^{13}$C NMR (126 MHz, DMSO) δ 156.65, 153.75, 150.76, 135.58, 116.56, 90.06, 75.66, 74.71, 73.89, 66.95, 27.30, 26.83, 25.69, 22.23, 19.97, 18.02, -4.59, -5.13. HRMS (ESI/Q-TOF) calcd for $C_{24}H_{44}N_5O_5Si_2^+$ $[M + H]^+$ 538.2875, found 538.2890.

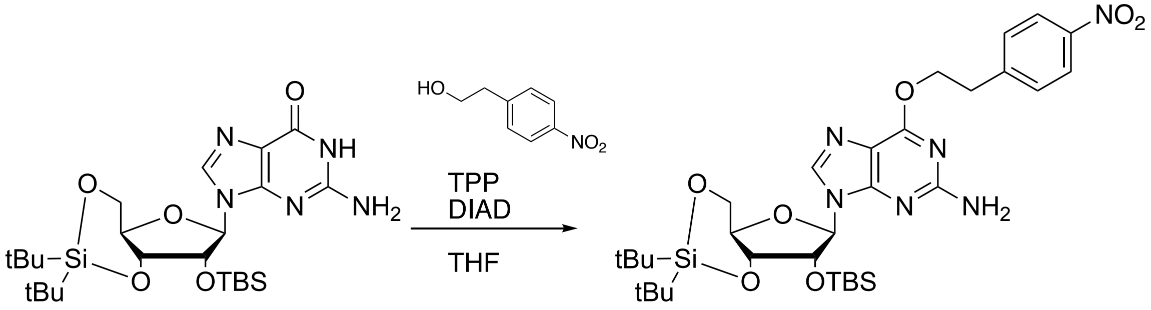

To a solution of triphenylphosphine (7.90 g, 30.1 mmol) and 4-nitrophenethanol (5.05 g, 30.1 mmol) in dioxane (80 mL) was added ribose-protected guanosine **13** (8.10 g, 15.1 mmol). The solution was heated to 100 °C and diisopropyl azodicarboxylate (DIAD, 5.91 mL, 30.1 mmol) was added dropwise. During the course of the addition of DIAD, the reaction mixture became homogenous. After 1h, the reaction was allowed to cool to 23 °C, and then solvent was removed under reduced pressure. The crude residue was purified by silica flash chromatography (25% to 75% ethyl acetate in hexanes) to yield *O*$^6$-(4-nitrophenethyl)-guanosine as a white solid (6.72 g, 65% yield). $^1$H NMR (500 MHz, $CDCl_3$) δ 8.19 – 8.12 (m, 2H), 7.66 (s, 1H), 7.52 – 7.45 (m, 2H), 5.79 (s, 1H), 4.85 (s, 2H), 4.73 (td, *J* = 6.8, 2.3 Hz, 2H), 4.50 (d, *J* = 4.6 Hz, 1H), 4.48 (dd, *J* = 9.2, 5.1 Hz, 1H), 4.43 (dd, *J* = 9.6, 4.7 Hz, 1H), 4.19 (td, *J* = 10.0, 5.1 Hz, 1H), 4.01 (dd, *J* = 10.5, 9.2 Hz, 1H), 3.27 (t, *J* = 6.8

Hz, 2H), 1.05 (d, *J* = 18.6 Hz, 19H), 0.93 (s, 9H), 0.15 (s, 3H), 0.14 (s, 3H). $^{13}$C NMR (126 MHz, CDCl$_3$) δ 160.78, 159.09, 153.23, 146.99, 145.98, 137.72, 130.09, 123.87, 115.83, 92.34, 75.96, 75.54, 74.70, 67.98, 66.67, 35.30, 27.60, 27.13, 26.02, 22.91, 20.46, 18.47, -4.13, -4.86. HRMS (ESI/Q-TOF) calcd for $C_{32}H_{51}N_6O_7Si_2^+$ [M + H]$^+$ 687.3352, found 687.3357.

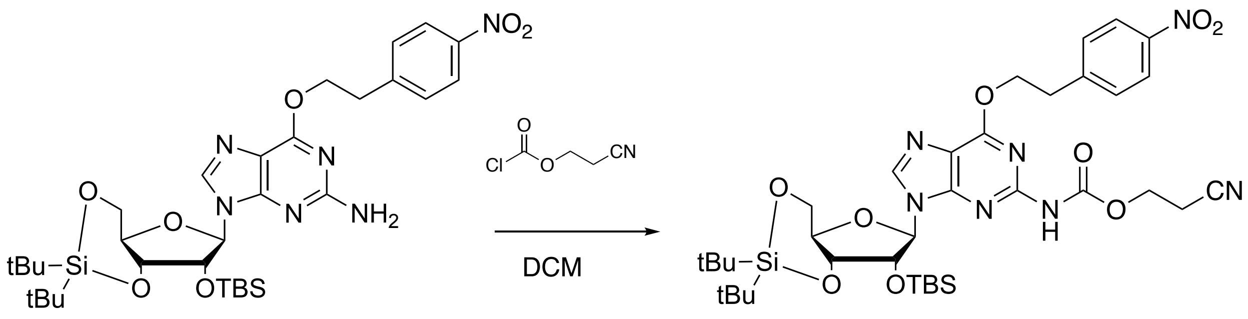


*O*$^6$-(4-nitrophenethyl)-guanosine (6.00 g, 8.74 mmol) was dissolved in anhydrous dichloromethane (50 mL) and ceoc-chloroformate (1.75 g, 13.11 mmol) was added dropwise. After 2 h of stirring at 23 °C, an addition portion of ceoc-chloroformate (0.800 g, 5.99 mmol) was added and the reaction was stirred for an additional 2 h. The reaction was then diluted with ethyl acetate (300 mL), transferred to a separatory funnel, and washed with water (3 x 100 mL) and brine (2 x 100 mL). The organic layer was then dried over anhydrous $Na_2SO_4$, filtered, and then condensed under reduced pressure. The crude product was purified by silica flash chromatography (33% to 50% ethyl acetate in hexanes) to yield **14** (5.13 g, 75% yield). $^1$H NMR (500 MHz, CDCl$_3$) δ 8.21 – 8.13 (m, 2H), 7.90 (s, 1H), 7.56 – 7.50 (m, 2H), 7.37 (s, 1H), 5.86 (s, 1H), 4.82 (td, *J* = 6.8, 1.3 Hz, 2H), 4.59 (d, *J* = 4.9 Hz, 1H), 4.50 – 4.35 (m, 4H), 4.22 – 4.09 (m, 1H), 4.03 (dd, *J* = 10.5, 9.2 Hz, 1H), 3.32 (t, *J* = 6.8 Hz, 2H), 2.78 (dt, *J* = 12.5, 6.3 Hz, 2H), 1.08 (s, 9H), 1.04 (s, 8H), 0.91 (s, 8H), 0.13 (d, *J* = 1.4 Hz, 6H). $^{13}$C NMR (126 MHz, CDCl$_3$) δ 160.76, 152.02, 151.71, 150.19, 147.02, 145.72, 140.10, 130.22, 123.89, 118.23, 116.86, 92.66, 75.96, 75.43, 74.85, 67.76, 67.32, 62.53, 59.77, 35.14, 27.59, 27.15, 26.00, 22.85, 20.49, 18.47, -4.14, -4.89. HRMS (ESI/Q-TOF) calcd for $C_{36}H_{53}N_7NaO_9Si_2^+$ [M + Na]$^+$ 806.3336, found 806.3343.

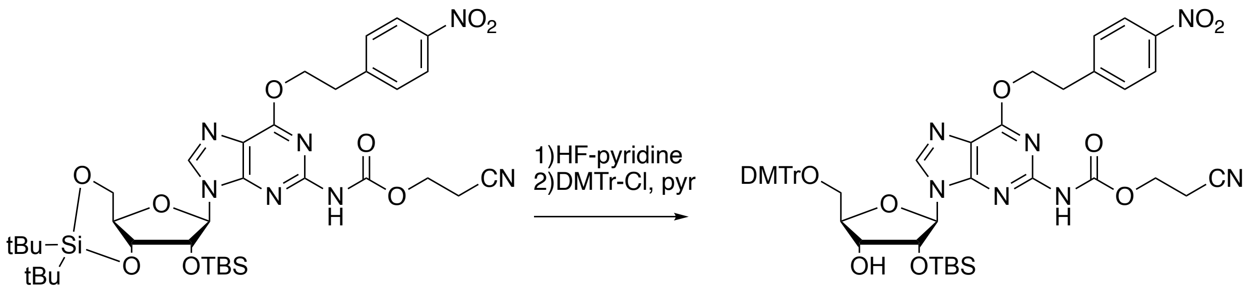


HF-pyridine (0.60 mL, 23.0 mmol) was added to pyridine (4.20 mL, 52.0 mmol) in a 15 mL plastic conical tube at 0 °C. Compound **14** (4.50 g, 5.74 mmol) was suspended in DCM (29 mL) in a 50 mL plastic conical tube and cooled to 0 °C. The HF-pyridine solution was added dropwise to the DCM suspension of **14** at 0 °C with stirring. After 90 minutes, the reaction mixture was diluted with DCM (150 mL), and washed with 150 mL each of water, saturated aqueous sodium bicarbonate, and brine. The organic layer was collected and dried over anhydrous $Na_2SO_4$, filtered, and then condensed under reduced pressure. The crude diol was dissolved in pyridine (11.4 mL) and cooled to 0 °C. Dimethoxytrityl chloride (2.14 g, 6.31 mmol) was added to the pyridine solution at 0 °C with stirring. After 10 minutes, the seal reaction vessel was transferred to a 4 °C refrigerator for

16 h. To quench the reaction, methanol (3 mL) was added, and the reaction mixture was stirred at 23 °C for 5 minutes after which, all volatiles were removed under reduced pressure. The crude product was purified by silica flash chromatography to yield **15** (4.33 g, 80% yield). $^1$H NMR (500 MHz, $CDCl_3$) δ 8.20 – 8.14 (m, 2H), 7.82 (s, 1H), 7.52 (d, *J* = 8.7 Hz, 2H), 7.47 – 7.42 (m, 1H), 7.35 – 7.22 (m, 3H), 7.20 – 7.14 (m, 3H), 6.86 – 6.80 (m, 3H), 5.73 (d, *J* = 7.2 Hz, 1H), 5.06 (dd, *J* = 7.2, 4.9 Hz, 1H), 4.95 (d, *J* = 11.0 Hz, 1H), 4.85 (qt, *J* = 10.8, 6.8 Hz, 2H), 4.47 – 4.35 (m, 3H), 4.32 (s, 1H), 4.12 (q, *J* = 7.1 Hz, 1H), 3.95 (d, *J* = 12.8 Hz, 1H), 3.80 (s, 5H), 3.80 – 3.72 (m, 1H), 3.33 (t, *J* = 6.9 Hz, 2H), 2.82 – 2.73 (m, 3H), 2.04 (s, 2H), 1.26 (t, *J* = 7.1 Hz, 2H), 0.86 – 0.79 (m, 1H), 0.81 (s, 6H), -0.13 (s, 2H), -0.35 (s, 2H). HRMS (ESI/Q-TOF) calcd for $C_{49}H_{56}N_7O_{11}Si^+$ $[M + H]^+$ 946.3802, found 946.3827.

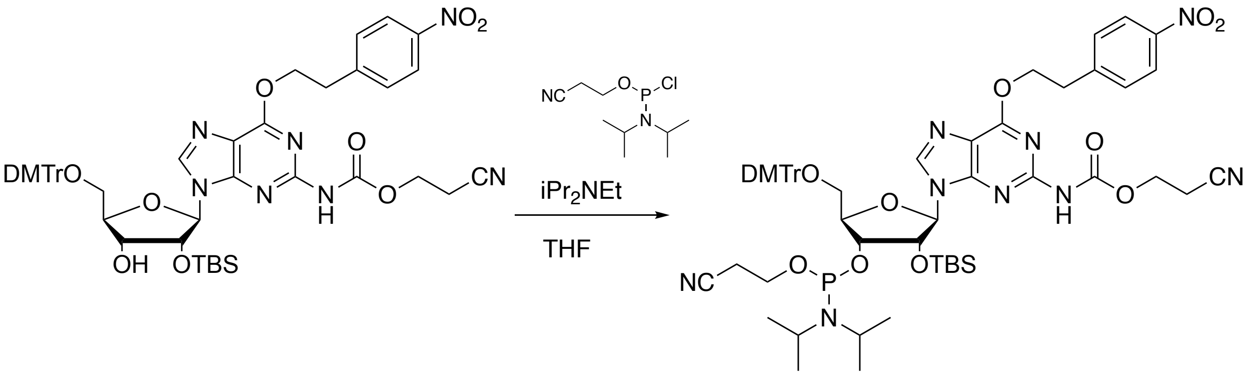


Free 3'OH-guanosine **15** (0.942 g, 1.00 mmol) was dissolved in THF (10 mL) and cooled to 0 °C. $iPr_2NEt$ (0.61 mL, 3.5 mmol) was added immediately followed by addition of 2-cyanoethyl *N,N*-diisopropyl phosphoamidochloridite (0.31 mL, 1.39 mmol). The reaction mixture was allowed to warm to 23 °C with stirring. After 5 h, the reaction was quenched by addition of MeOH (1 mL) and stirred an additional 30 min at 23 °C. The reaction mixture was then diluted with ethyl acetate (50 mL), transferred to a separatory funnel, and washed with sat'd aq $NaHCO_3$ (3 x 50 mL). The organic layer was then dried over anhydrous $Na_2SO_4$, filtered, and then condensed under reduced pressure. Purification by silica flash chromatography yield *N*-protected G phosphoramidite **16** (0.692 g, 60% yield). $^1$H NMR (500 MHz, $CDCl_3$) δ 8.18 – 8.13 (m, 2H), 8.03 (d, *J* = 10.0 Hz, 1H), 7.53 (dd, *J* = 8.7, 2.2 Hz, 2H), 7.51 – 7.48 (m, 1H), 7.48 – 7.44 (m, 1H), 7.43 (s, 1H), 7.40 – 7.31 (m, 6H), 7.30 – 7.23 (m, 1H), 7.24 – 7.18 (m, 1H), 6.85 – 6.72 (m, 5H), 6.01 (d, *J* = 7.2 Hz, 1H), 5.87 (d, *J* = 6.9 Hz, 0H), 5.12 (dd, *J* = 7.0, 5.1 Hz, 0H), 5.00 (dd, *J* = 7.2, 4.6 Hz, 1H), 4.85 (td, *J* = 6.9, 2.3 Hz, 2H), 4.42 – 4.39 (m, 0H), 4.38 – 4.33 (m, 1H), 4.32 – 4.28 (m, 1H), 4.27 – 4.25 (m, 1H), 4.23 (t, *J* = 6.3 Hz, 1H), 4.09 – 4.00 (m, 1H), 3.97 – 3.87 (m, 1H), 3.80 – 3.72 (m, 7H), 3.67 – 3.53 (m, 3H), 3.49 (dd, *J* = 10.7, 3.0 Hz, 1H), 3.34 (td, *J* = 6.9, 1.9 Hz, 2H), 3.31 – 3.23 (m, 1H), 2.72 (td, *J* = 6.7, 3.3 Hz, 1H), 2.68 (t, *J* = 6.3 Hz, 2H), 2.61 (td, *J* = 6.3, 2.2 Hz, 1H), 2.25 (qt, *J* = 16.7, 6.6 Hz, 1H), 1.40 – 1.34 (m, 1H), 1.33 – 1.23 (m, 1H), 1.22 – 1.14 (m, 8H), 1.01 (d, *J* = 6.8 Hz, 4H), 0.75 (s, 9H), -0.01 (s, 1H), -0.06 (s, 2H), -0.25 (d, *J* = 4.2 Hz, 3H). $^{13}$C NMR (126 MHz, $CDCl_3$) δ 160.78, 160.74, 158.73, 158.70, 153.28, 151.59, 150.49, 150.41, 146.96, 145.89, 144.59, 141.40, 140.59, 135.62, 135.56, 130.21 (d, *J* = 2.9 Hz), 130.12, 130.10, 128.44, 128.29, 128.13, 128.09, 128.03, 127.15, 123.83, 118.54, 117.96, 116.88, 113.38, 113.29, 88.39, 87.42, 86.85, 84.69, 84.33, 84.30, 75.79, 74.00, 72.68, 72.57, 67.05, 67.01, 63.57, 63.44, 59.49, 59.44, 59.22, 59.10, 55.36, 55.34, 43.59, 43.49, 43.03, 42.93, 35.19, 25.72, 25.70, 25.66, 24.90, 24.84, 24.77, 24.74, 24.69, 20.54, 20.50, 18.28, 18.20, 18.06, 17.94, -4.61, -4.64, -5.09.

*Photocleavable linker*

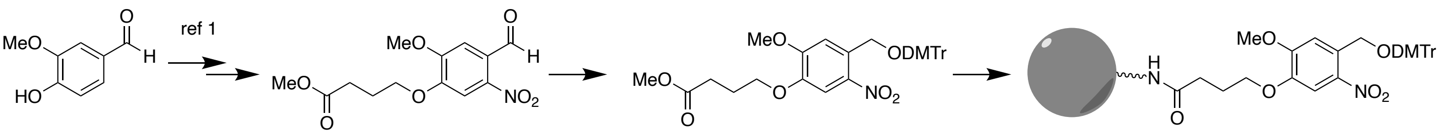


**Vanillin** **17** **18** **19**

**Scheme S5.** Synthesis of photocleavable solid support.

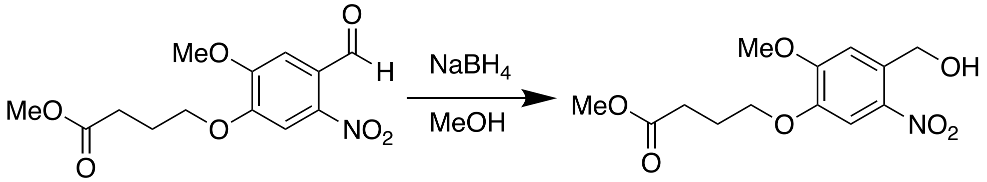


Aldehyde **17** was synthesized in two steps from vanillin as described by Andriollo et al. (2019). To an ice cold solution of aryl aldehyde (5.77g, 19.4 mmol) in 1:1 THF:MeOH (74 mL) was added sodium borohydride (1.1 g, 29.1 mmol) in portions. After 15 min at 0 °C, the reaction was complete by TLC (6:4 hexanes:ethyl acetate). Solvent was removed under reduced pressure and the resulting residue was quenched with aq. $NH_4Cl$ (50 mL). The aqueous layer was extracted with DCM (3 x 30 mL). The DCM layer was then washed with brine (50 mL) and dried over anhydrous $Na_2SO_4$, filtered, and then condensed under reduced pressure to yield nitroveratrol **18** (5.38 g, 93% yield) without further purification. $^1$H NMR (500 MHz, $CDCl_3$) δ 7.67 (s, 1H), 7.16 (s, 1H), 4.93 (s, 2H), 4.11 (t, *J* = 6.3 Hz, 2H), 3.96 (s, 3H), 3.69 (s, 3H), 2.55 (t, *J* = 7.2 Hz, 2H), 2.18 (quint, *J* = 6.8 Hz, 2H). $^{13}$C NMR (126 MHz, $CDCl_3$) δ 173.51, 154.40, 147.23, 139.66, 132.59, 111.19, 109.61, 68.37, 62.84, 56.49, 51.85, 30.47, 24.36. HRMS (ESI/Q-TOF) calcd for $C_{13}H_{17}NNaO_7^+$ $[M + Na]^+$ 322.0897, found 322.0899.

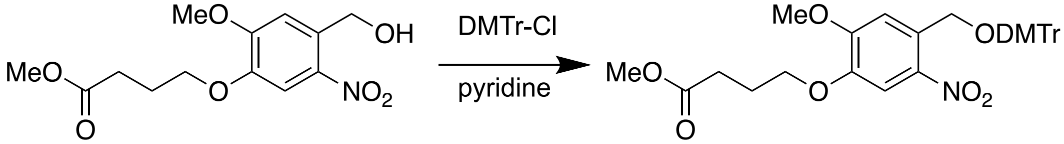


To an ice-cold solution of nitroveratrol **18** (4.20 g, 14.0 mmol) in DCM (46 mL) was added $iPr_2NEt$ (12.2 mL, 70 mmol) and DMTr-Cl (5.70 g, 16.8 mmol). After 30 min at 0 °C, the reaction mixture was allowed to warm to 23 °C. After 1 h at 23 °C, the reaction was complete by TLC (3:1 hexanes:ethyl acetate). The reaction mixture was diluted with DCM (100 mL), transferred to a separatory funnel, and washed with aq. $NaHCO_3$ (sat'd, 100 mL). The DCM solution was dried over anhydrous $Na_2SO_4$, filtered, and then condensed under reduced pressure. Purification by silica flash chromatography (0 to 50% ethyl acetate in hexanes with 1 % triethylamine) yielded pure DMTr-protected nitroveratrol product **19** (5.513 g, 65% yield). $^1$H NMR (500 MHz, $CDCl_3$) δ 7.75 (d, *J* = 4.0 Hz, 2H), 7.58 (d, *J* = 7.5 Hz, 2H), 7.46 (d, *J* = 9.0 Hz, 4H), 7.39 (t, *J* = 7.7 Hz, 2H), 7.36 – 7.30 (m, 1H), 7.29 – 7.24 (m, 0H), 6.93 (d, *J* = 9.0 Hz, 4H), 4.76 (s, 2H), 4.20 (t, *J* = 6.2 Hz, 2H), 4.12 (s, 3H), 3.88 (s, 6H), 3.79 (s, 3H), 2.65 (t, *J* = 7.2 Hz, 2H), 2.28 (quint, *J* = 6.8 Hz, 2H). $^{13}$C NMR (126 MHz, $CDCl_3$) δ 173.48, 158.75, 154.19, 146.51, 144.91, 138.91, 135.93, 132.18, 130.10, 129.24, 128.14, 128.08, 127.94, 127.87, 127.11, 113.37, 111.23, 109.93, 109.36, 87.23, 68.33, 63.17, 56.34, 55.33, 51.82, 30.50, 24.40. HRMS (ESI/Q-TOF) calcd for $C_{34}H_{35}NNaO_9^+$ [M + Na]+ 624.2204, found 624.2198.

***Preparation of photolabile CPG resin***

DMTr-protected photolabile linker **19** (0.020 g, 0.033 mmol) was dissolved in THF (0.1 mL) and aq. LiOH (2M, 0.018 mL, 0.036 mmol) was added. After stirring overnight at 23 °C, solvent was removed under reduced pressure. The resulting residue was coevaporated twice with anhydrous acetonitrile to minimize water content. The residue was suspended in acetonitrile (0.4 mL) and $iPr_2NEt$ (0.017 mL, 0.049 mmol) was added followed by HATU (0.0125 g, 0.033 mmol). After stirring for 5 min, the reaction mixture became homogenous at which point, it was diluted with anhydrous acetonitrile (4.4 mL) and the stir bar was removed. CPG-LCAA (0.500 g, 132 μmol amine/g) was added and the reaction was manually swirled to mix. After 5 min, the reaction mixture was manually swirled again, this process was repeated for a total of 20 min of reaction time. The solid support was then filtered and washed with 10 mL each of dichloromethane, acetonitrile, water, acetonitrile, and dichloromethane. After drying *in vacuo* for 1 h, free amines on the solid support were capped by addition of a solution of 2,6-lutidine (0.25 mL), pivaloyl chloride (0.25 mL), and N-methyl imidazole (0.25 mL) in THF (4.25 mL). After overnight rocking, the solid support was washed again with 10 mL each of dichloromethane, acetonitrile, water, acetonitrile, and dichloromethane. After drying *in vacuo*, for 1 h, loading was assessed by trityl assay.

***Solid-phase RNA oligonucleotide synthesis***

All RNA syntheses were performed on a 1 μmol scale using *N*-ceoc-protected phosphoramidites of adenosine and cytosine, *N*-unprotected guanosine, dihydrouridine phosphoramidite, and commercial *N*-acetylated cytosine and uridine. All oligonucleotides were synthesized on CPG-solid support derivatized with a photocleavable universal linker. All phosphoramidites were prepared as 0.1 M solutions in anhydrous acetonitrile except for *N*-unprotected guanosine phosphoramidite which was prepared as a 0.1 M solution in anhydrous dichloromethane.

| Reagent | Volume | Time |
|---|---|---|
| 3% TCA in DCM | 1.5 mL | 90 s |
| Acetonitrile | 3.0 mL | 50 s |
| ETT (0.25 M in acetonitrile) | 0.45 mL | 360 s |
| Phosphoramidite | 0.20 mL | 360 s |
| ETT (first coupling only) | 0.45 mL | 360 s |
| Phosphoramidite (first coupling only) | 0.20 mL | 360 s |
| Acetonitrile | 0.60 mL | 10 s |
| $I_2$ (0.02 M in 7:2:1 THF/pyridine/$H_2O$) | 0.60 mL | 50 s |
| Acetonitrile | 3.0 mL | 50 s |

***Deprotection and cleavage of RNA oligonucleotides***

*On-column deprotection*

Deprotection was initiated immediately after oligonucleotide synthesis. Automated DBU-mediated removal of *N*-ceoc and cyanoethyl phosphate protecting groups was performed "on-column" using the following method.

| Reagent | Volume | Time |
|---|---|---|
| Acetonitrile | 5.0 mL | 50 s |
| DBU (0.5M in acetonitrile) | 3.5 mL | 5 min |
| Acetonitrile | 5.0 mL | 50 s |
| DBU (0.5M in acetonitrile) | 3.5 mL | 5 min |
| Acetonitrile | 5.0 mL | 50 s |
| DBU (0.5M in acetonitrile) | 2.0 mL | 1 min |
| DBU (0.5M in acetonitrile) | 5.0 mL | 60 min |
| Acetonitrile | 2.0 mL | 10 s |
| DBU (0.5M in acetonitrile) | 2.0 mL | 1 min |
| DBU (0.5M in acetonitrile) | 5.0 mL | 60 min |
| Acetonitrile | 2.0 mL | 10 s |
| DBU (0.5M in acetonitrile) | 2.0 mL | 1 min |
| DBU (0.5M in acetonitrile) | 2.0 mL | 20 min |
| Acetonitrile | 12.0 mL | 3 min |

After *N*-ceoc removal, 5'-ODMTr groups were removed by passing 3% TCA in DCM through the column for 1.5 min. After rinsing the column with acetonitrile for 1 min, it was purged with argon, and then dried *in vacuo* for 10 min.

*Photolysis*

Solid support was suspended in acetonitrile:TEAA buffer (100 mM, pH 7) (9:1, 2 mL) in a 2 mL glass HPLC vial. The vial was taped to a 365 nm LED light and irradiated while rocking for 1 h. The solvent was passed through a 0.45 μm PTFE filter to remove any solids and the solution was lyophilized to dryness.

*2'-OTBS removal*

Crude RNA from photolysis was dissolved in DMSO (100 μL), mixed by vortexing, and anhydrous $iPr_2NEt$ (70 μL) was added followed by $Et_3N$-3HF (62.5 μL). The solution was mixed by vortex then incubated at 65 °C for 2.5 h. Desilylated RNA was incubated at -20 °C for 2 min. NaOAc (25 μL, 3.0 M, pH 5.5) was added to the DMSO solution followed by dilution in *n*-BuOH (1 mL). The mixture was incubated at -20 °C for 30 minutes then centrifuged at 12500 X g for 10 min at 4 °C. Solvent was aspirated carefully. The pellet was then washed with ethanol (0.7 mL) twice to remove any residual *n*-BuOH. Finally, the pellet was dried *in vacuo* for 10 min to removal any residual ethanol.

***Purification of RNA oligonucleotides***

Crude oligonucleotides were purified by polyacrylamide gel electrophoresis (PAGE) using a 20% acrylamide gel. RNA was visualized by UV-shadowing using a TLC plate illuminated with 254 nm light. Desired bands were excised, crushed, and soaked in buffer (0.5 M $NH_4OAc$, 0.02M EDTA) for 16 h at 4 °C. Solutions containing purified RNA was separated from gel using a spin filter. Solutions were lyophilized to dryness.

Pure RNA was desalted using $C_{18}$ reversed phase cartridges. Briefly, the $C_{18}$cartridge was conditioned with acetonitrile (10 mL), acetonitrile:NaOAc (100 mM) (1:1, 10 mL), and NaOAc (100 mM, 10 mL). RNA was dissolved in NaOAc (100 mM, 5 mL) and loaded onto the column. The column was then washed with TEAA (10 mM, 10 mL). RNA was eluted in 1 mL fractions with acetonitrile:TEAA (10 mM) (7:3, 5 mL). Fractions containing RNA, as determined by absorbance at 260 nm, were pooled and lyophilized to yield pure desalted RNA. $tRNA^{Ser}$ D-loop containing $ac^4C$ required further purification by HPLC. Flow rate 1 mL/min; solvent A: aq. TEAA (20 mM, pH 6.8); solvent B: acetonitrile. Gradient: 3% B 0 – 2 min, 3% to 11.6% B 2 – 15 min, 11.6% B to 95% B 15 – 16 min, 95% B 16 – 19 min.

**UV Melting Analysis**

***Hybridization of duplex RNAs***

Custom RNA oligonucleotide top strand containing ac4C was synthesized, PAGE-purified, and desalted prior to hybridization for thermal denaturation studies. PAGE-purified top strand without ac4C was purchased from Integrated DNA Technologies (Coralville, IA) with intact 3' phosphate. Complementary RNA bottom strands were purchased from Integrated DNA Technologies (Coralville, IA) at their highest purity, with the following sequence.

| | |
|---|---|
| Top Strand C: | 5'-CUUCCGUAGGp-3' |
| Match bottom strand: | 5'-CCUACGGAAG-3' |
| A Mismatch bottom strand: | 5'-CCUAC**A**GAAG-3' |
| C Mismatch bottom strand: | 5'-CCUAC**C**GAAG-3' |
| U Mismatch bottom strand: | 5'-CCUAC**U**GAAG-3' |
| GU in bottom strand: | 5'-CCUA**U**GGAAG-3' |
| GU out bottom strand: | 5'-CCU**G**CGGAAG-3' |
| GA mismatch bottom strand: | 5'-CCUA**A**GGAAG-3' |

Prior to thermal denaturation, strands were suspended 1X $T_m$ buffer (1 M NaCl, 10 mM sodium phosphate [pH 7.4]) with complementary bottom strands. The final equimolar concentration of each RNA strand was 10 μM. To facilitate hybridization, duplexes were heated to 95 °C for 5 minutes and slowly cooled to room temperature over a period of 1 hour. Duplexes were stored at -80 ˚C prior to UV melting temperature determination.

***Thermal denaturation of RNA duplexes***

Absorbance versus temperature profiles of RNA duplexes were recorded at 260 nm on a PerkinElmer Photodiode Array Lambda 465 UV/Vis Spectrophotometer equipped with Starna Type 26/LHS. Sub-micro, low head space (26.50/LHSQ10) cuvettes and a Peltier temperature control device. For melting temperature determinations, each RNA duplex was diluted the desired concentration in 1X $T_m$ buffer (1 M NaCl, 10 mM sodium phosphate [pH 7.4]). 75 μL of each sample was used for analysis. Melting profiles were collected over a temperature range of 20 °C to 90 °C at a rate of 1 °C/min. All thermal denaturation experiments were performed in triplicate. Melting temperatures and thermodynamic parameters were calculated using Meltwin v3.5 software.[6,7]

***Annealing of hairpin RNA***

Custom RNA oligonucleotide hairpins with and without ac4C were synthesized, PAGE-purified, and HPLC-purified (for the $ac^4C$-containing hairpin only) and suspended 1X $T_m$ buffer (1 M NaCl, 10 mM sodium phosphate [pH 7.4]) at a final concentration of 10 μM. Hairpins were annealed by heating to 95 °C for 5 minutes and slowly cooled to room temperature over a period of 1 hour.

***Thermal denaturation of RNA hairpins***

Absorbance versus temperature profiles of RNA hairpins were recorded at 280 nm on a PerkinElmer Photodiode Array Lambda 465 UV/Vis Spectrophotometer equipped with Starna Type 26/LHS. Sub-micro, low head space (26.50/LHSQ10) cuvettes and a Peltier temperature control device. 75 μL of each sample was used for analysis. Melting profiles were collected over a temperature range of 20 °C to 95 °C at a rate of 1 °C/min. All thermal denaturation experiments were performed in triplicate. Melting temperatures and thermodynamic parameters were calculated using Meltwin v3.5 software.[6,7]

### UV melting curves and van't Hoff plots

ac4C-G

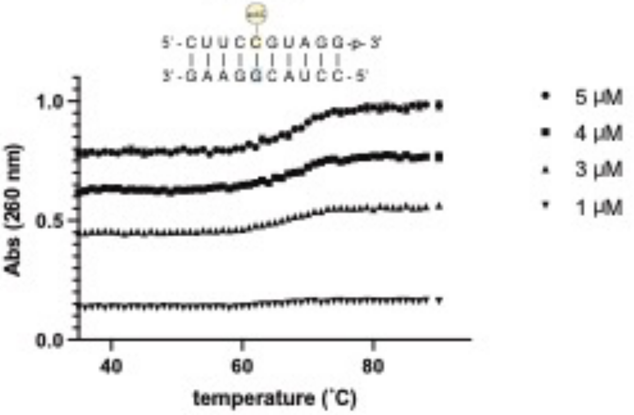


ac4C-A

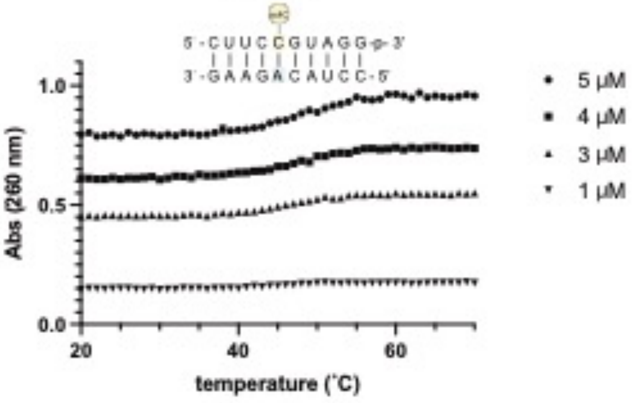


ac4C-U

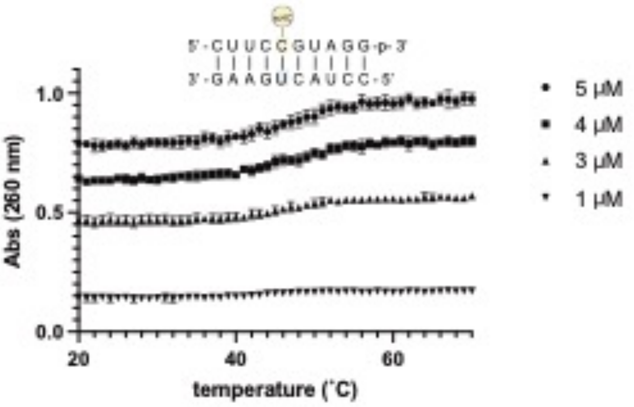


ac4C-C

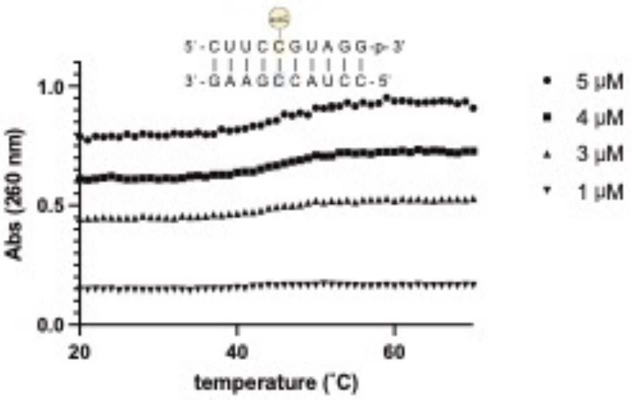


C-G

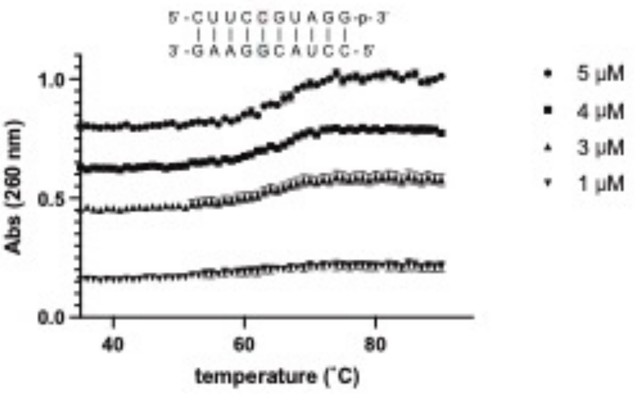


C-A

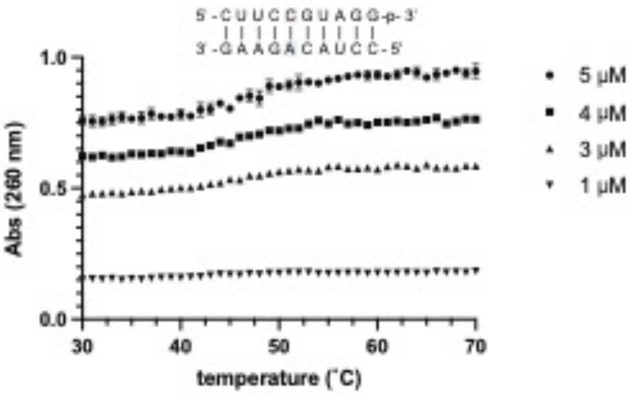


C-U

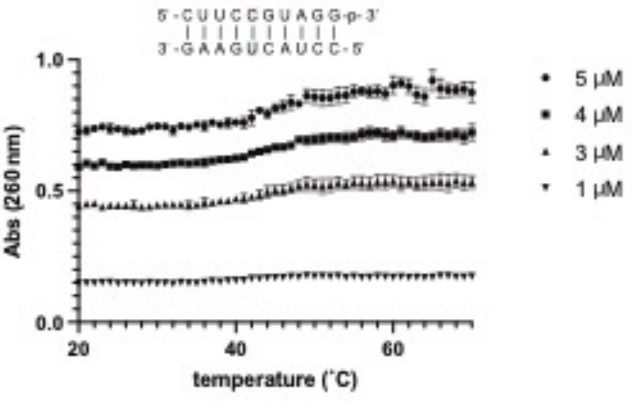


C-C

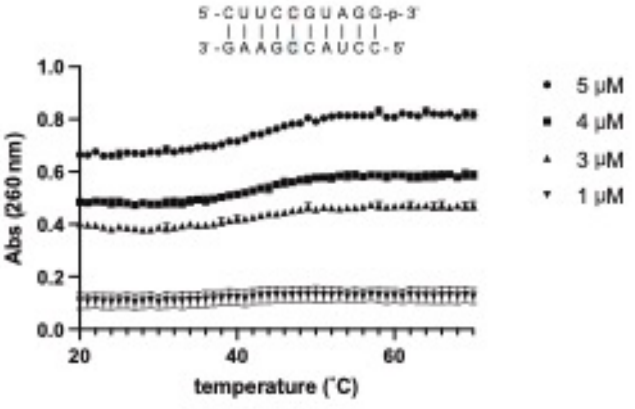

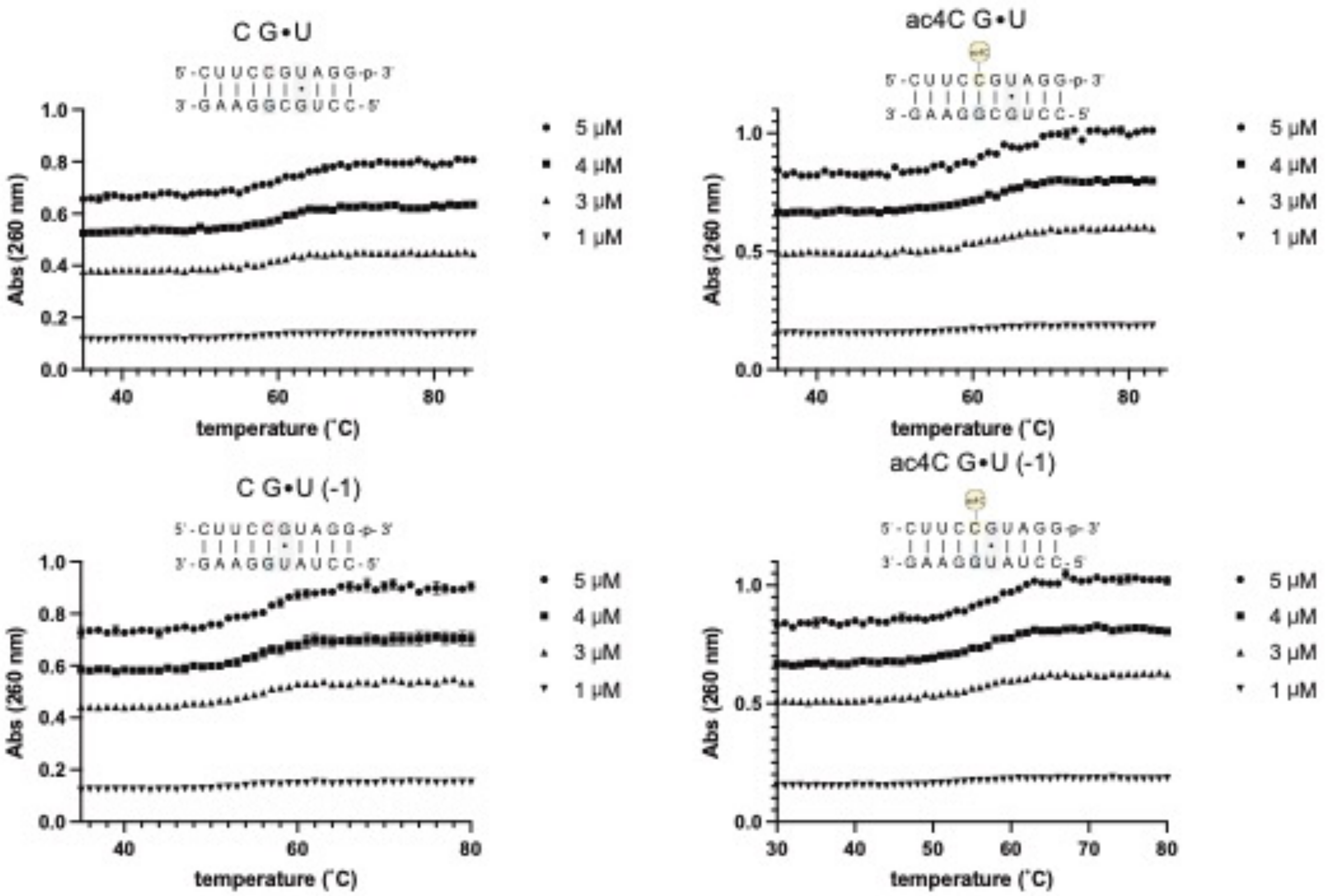
C G•U
5'-CUUCCGUAGG-p-3'
3'-GAAGGCGUCC-5'
ac4C G•U
5'-CUUCCGUAGG-p-3'
3'-GAAGGCGUCC-5'
C G•U (-1)
5'-CUUCCGUAGG-p-3'
3'-GAAGGUAUCC-5'
ac4C G•U (-1)
5'-CUUCCGUAGG-p-3'
3'-GAAGGUAUCC-5'
Abs (260 nm)
temperature (˚C)
5 µM
4 µM
3 µM
1 µM

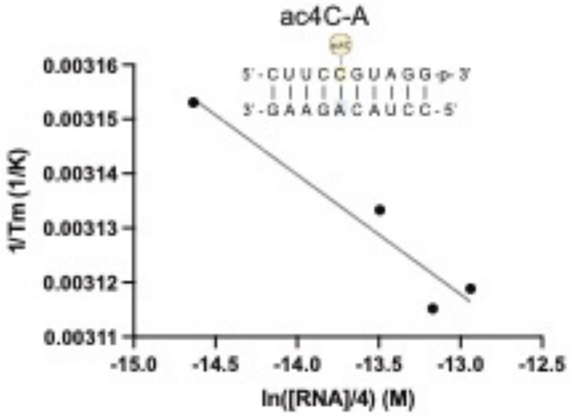
ac4C-A
5'-CUUCCGUAGG-p-3'
3'-GAAGACAUCC-5'
1/Tm (1/K)
0.00316
0.00315
0.00314
0.00313
0.00312
0.00311
-15.0
-14.5
-14.0
-13.5
-13.0
-12.5
ln([RNA]/4) (M)

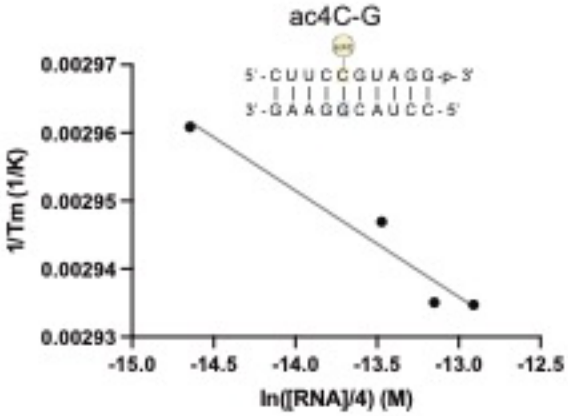
ac4C-G
5'-CUUCCGUAGG-p-3'
3'-GAAGGCAUCC-5'
1/Tm (1/K)
0.00297
0.00296
0.00295
0.00294
0.00293
-15.0
-14.5
-14.0
-13.5
-13.0
-12.5
ln([RNA]/4) (M)

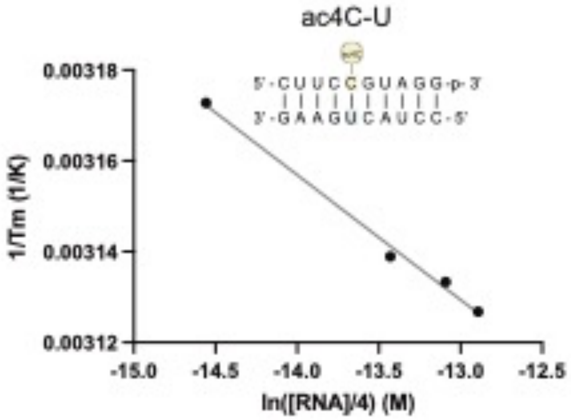
ac4C-U
5'-CUUCCGUAGG-p-3'
3'-GAAGUCAUCC-5'
1/Tm (1/K)
0.00318
0.00316
0.00314
0.00312
-15.0
-14.5
-14.0
-13.5
-13.0
-12.5
ln([RNA]/4) (M)

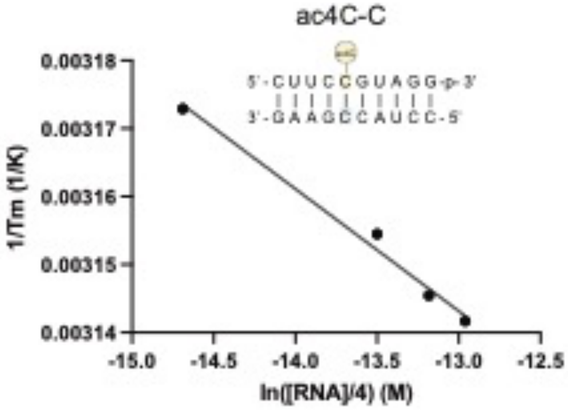
ac4C-C
5'-CUUCCGUAGG-p-3'
3'-GAAGCCAUCC-5'
1/Tm (1/K)
0.00318
0.00317
0.00316
0.00315
0.00314
-15.0
-14.5
-14.0
-13.5
-13.0
-12.5
ln([RNA]/4) (M)

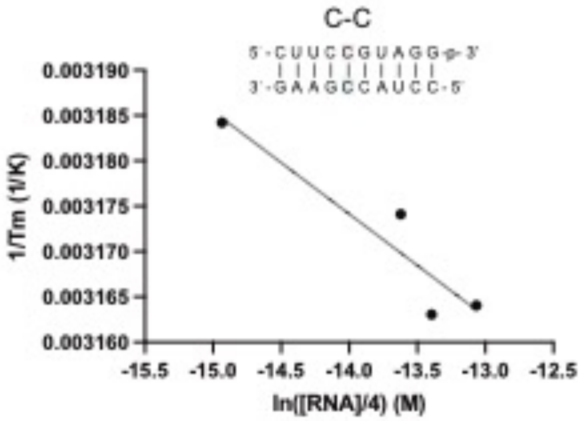
C-C
5'-CUUCCGUAGG-p-3'
3'-GAAGCCAUCC-5'
1/Tm (1/K)
0.003190
0.003185
0.003180
0.003175
0.003170
0.003165
0.003160
-15.5
-15.0
-14.5
-14.0
-13.5
-13.0
-12.5
ln([RNA]/4) (M)

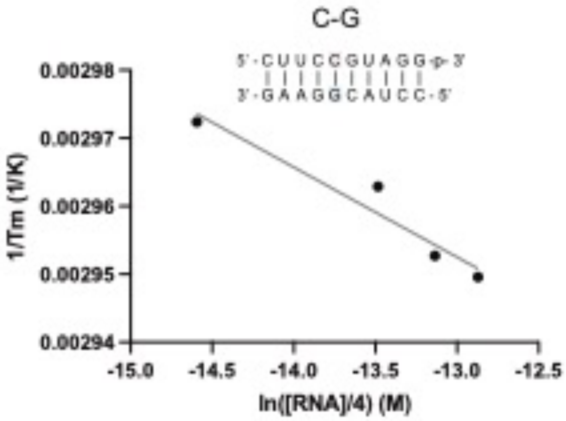
C-G
5'-CUUCCGUAGG-p-3'
3'-GAAGGCAUCC-5'
1/Tm (1/K)
0.00298
0.00297
0.00296
0.00295
0.00294
-15.0
-14.5
-14.0
-13.5
-13.0
-12.5
ln([RNA]/4) (M)

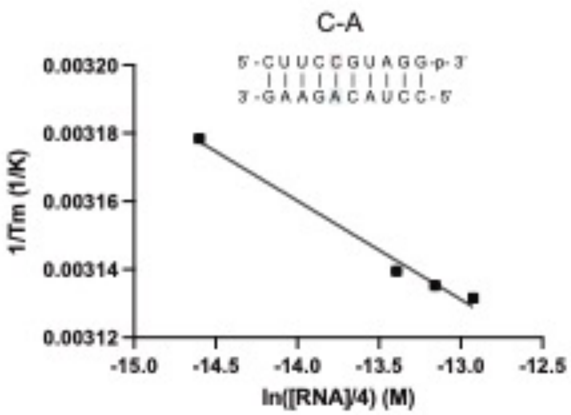
C-A
5'-CUUCCGUAGG-p-3'
3'-GAAGACAUCC-5'
1/Tm (1/K)
0.00320
0.00318
0.00316
0.00314
0.00312
-15.0
-14.5
-14.0
-13.5
-13.0
-12.5
ln([RNA]/4) (M)

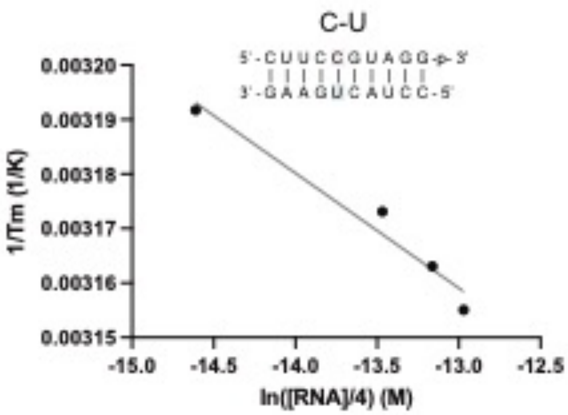
C-U
5'-CUUCCGUAGG-p-3'
3'-GAAGUCAUCC-5'
1/Tm (1/K)
0.00320
0.00319
0.00318
0.00317
0.00316
0.00315
-15.0
-14.5
-14.0
-13.5
-13.0
-12.5
ln([RNA]/4) (M)

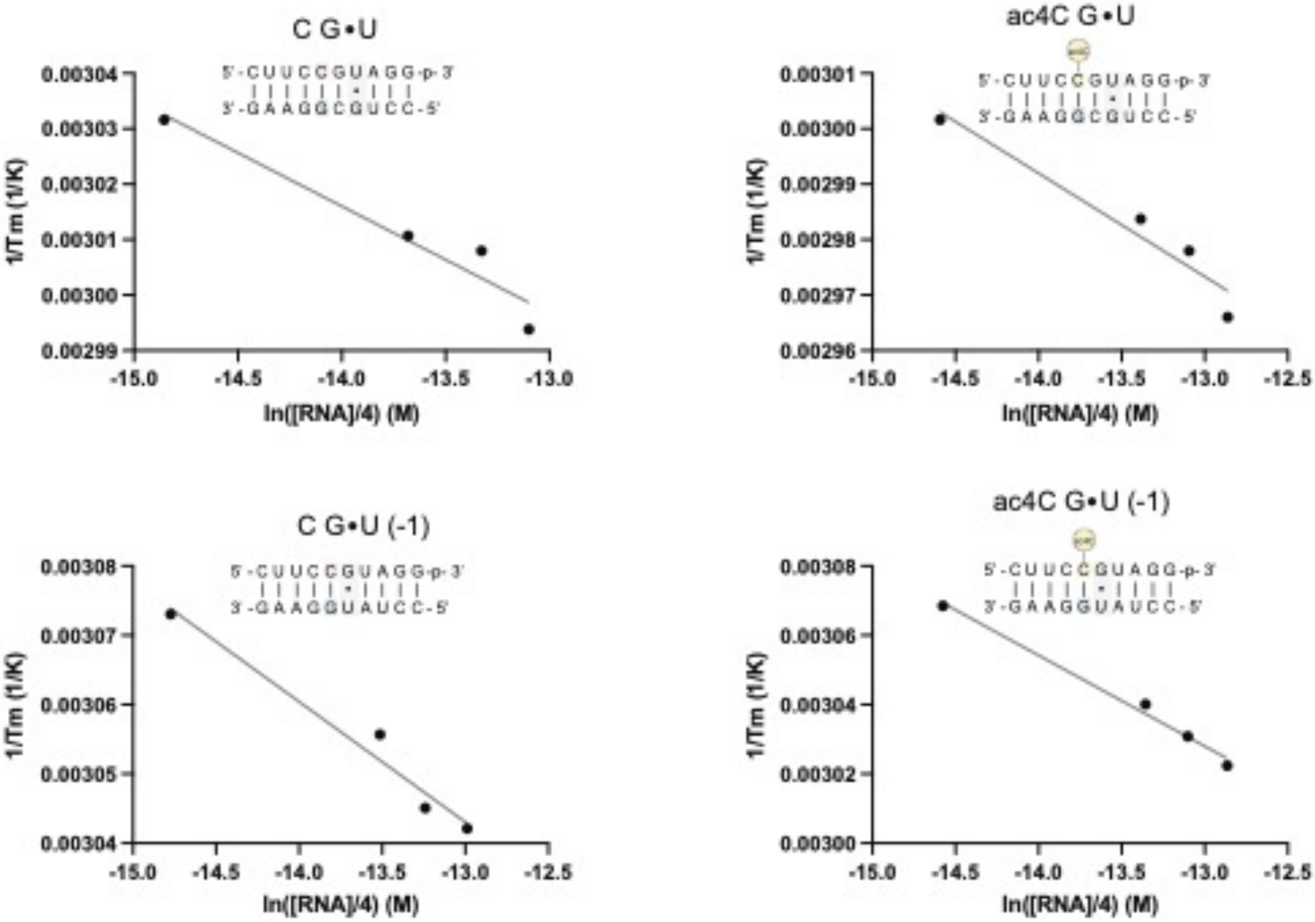
C G•U
5'-CUUCCGUAGG-p-3'
3'-GAAGGCGUCC-5'
1/Tm (1/K)
ln([RNA]/4) (M)
ac4C G•U
5'-CUUCCGUAGG-p-3'
3'-GAAGGCGUCC-5'
1/Tm (1/K)
ln([RNA]/4) (M)
C G•U (-1)
5'-CUUCCGUAGG-p-3'
3'-GAAGGUAUCC-5'
1/Tm (1/K)
ln([RNA]/4) (M)
ac4C G•U (-1)
5'-CUUCCGUAGG-p-3'
3'-GAAGGUAUCC-5'
1/Tm (1/K)
ln([RNA]/4) (M)

**References:**


1. Xu, J., Duffy, C. D., Chan, C. K. W. & Sutherland, J. D. Solid-phase synthesis and hybrization behavior of partially 2′/3′-O-acetylated RNA oligonucleotides. *J. Org. Chem.* **79**, 3311–3326 (2014).
2. Sas-Chen, A. *et al.* Dynamic RNA acetylation revealed by quantitative cross-evolutionary mapping. *Nature* **583**, 638–643 (2020).
3. Stern, L. & Schulman, L. H. The role of the minor base N4-acetylcytidine in the function of the Escherichia coli noninitiator methionine transfer RNA. *J. Biol. Chem.* **253**, 6132–6139 (1978).
4. E. Gottlieb, H., Kotlyar, V. & Nudelman, A. NMR Chemical Shifts of Common Laboratory Solvents as Trace Impurities. *J. Org. Chem.* **62**, 7512–7515 (1997).
5. Andriollo, P. *et al.* C8-Linked Pyrrolobenzodiazepine Monomers with Inverted Building Blocks Show Selective Activity against Multidrug Resistant Gram-Positive Bacteria. *ACS Infect. Dis.* **4**, 158–174 (2018).
6. McDowell, J. A. & Turner, D. H. Investigation of the structural basis for thermodynamic stabilities of tandem GU mismatches: Solution structure of (rGAGGUCUC)2 by two-dimensional NMR and simulated annealing. *Biochemistry* **35**, 14077–14089 (1996).
7. Schroeder, S. J. & Turner, D. H. Optical melting measurements of nucleic acid thermodynamics. *Methods Enzymol.* **468**, 371–387 (2009).
